# Developing a Novel *In Vitro* Toxicity Assay for Predicting Inhalation Toxicity in Rats

**DOI:** 10.1101/2025.07.31.668016

**Authors:** Arpamas Vachiraarunwong, Masaki Fujioka, David B. Alexander, Shugo Suzuki, Runjie Guo, Guiyu Qiu, Yurina Kawamura, Kana Shibano, Ikue Noura, Kwanchanok Praseatsook, Hirotoshi Akane, Shinji Takasu, Hiroyuki Tsuda, Kumiko Ogawa, Hideki Wanibuchi, Min Gi

## Abstract

The development of alternative *in vitro* methods for assessing acute inhalation toxicity is a critical step toward reducing animal testing and aligns with the principles of the 3Rs (replacement, reduction, and refinement). In this study, we developed and optimized a neutral red uptake (NRU) assay using human lung adenocarcinoma cells (A549) as a predictive model (A549-NRU) for acute inhalation toxicity. To improve assay efficiency and robustness, we introduced two key modifications: the incubation time was reduced to 15 minutes to enable rapid and high-throughput screening, and for chemicals reactive with polystyrene 6-well glass plates were used to prevent chemical-induced degradation and ensure assay consistency. LC_50_ values were determined for 49 chemicals and compared with reported LC_50_ values from 4-hour rat inhalation studies. A significant positive correlation was observed between A549-NRU-derived LC_50_ values and *in vivo* LC_50_ values for water-soluble compounds and chemicals containing aldehyde, ketone, alcohol, ether, and epoxide functional groups, suggesting that *in vivo* LC_50_ values may be predictable using the A549-NRU assay. Additionally, A549-NRU LC_50_ values showed significant negative correlations with molecular weight and octanol–water partition coefficients, indicating that chemicals with higher values tended to be less cytotoxic *in vitro*. Importantly, the A549-NRU assay demonstrated stronger correlation with *in vivo* LC_50_ values than the conventional NRU assay using mouse 3T3 fibroblast cells. These findings support the use of the A549-NRU assay to estimate starting doses for *in vivo* studies, and potentially as an *in vitro* alternative for predicting acute inhalation toxicity.

**Impact statement:** The optimized A549-NRU assay demonstrates predictive potential for inhalation toxicity while reducing reliance on animal testing. This model serves as a human-relevant alternative for estimating starting doses for *in vivo* inhalation toxicity studies.

## Introduction

The toxicity assessment of hazardous chemicals is a critical concern in environmental and public health. Traditionally, the toxicity of hazardous chemicals has been determined using either the median lethal concentration (LC_50_) or median lethal dose (LD_50_), depending on the exposure route (OECD 2002; 2017; 2024). These parameters serve as benchmarks for classifying chemicals under the Globally Harmonized System of Classification and Labelling of Chemicals (GHS). These values are obtained from acute toxicity tests in animals, which estimate the concentration or amount of chemical that causes death in 50% of the test animals. Rodents are frequently used under controlled conditions (Kennedy and Jay Graepel 1991). Inhalation is a major route of human exposure to chemicals, requiring specific assessment using experimental animals to assess the risks of airborne substances (Bakand et al. 2005). However, standard whole-body acute inhalation toxicity studies (4-hour rat inhalation) according to OECD 403 require large and complex exposure equipment (OECD 2024), limiting implementation of these studies.

In addition, increasing concerns about animal welfare and ethical issues have led to the development of alternative approaches that align with the 3Rs principle, replacement, reduction, and refinement (Burden et al. 2015). Among these, *in vitro* cytotoxicity assays have emerged as practical tools for estimating chemical toxicity while reducing reliance on animal testing. In particular, *in vitro* to *in vivo* extrapolation (IVIVE) is increasingly applied in pharmacokinetics, drug metabolism, and toxicology to enhance the efficiency of hazard assessment and provide insights into human health risk prediction. In addition, IVIVE has also been used to estimate animal toxicity values, such as LC_50_ or LD_50_ to support dose selection and reduce the number of animals used in acute toxicity studies (Bell et al. 2018; Coecke et al. 2013; Krewski et al. 2020; OECD 2010).

The OECD Guidance Document 129 has adopted the neutral red uptake (NRU) assay, which uses mouse fibroblast 3T3 cells, for estimating starting doses for acute oral toxicity tests (OECD 2010). This assay assesses cell viability by measuring the uptake of neutral red dye, which penetrates cell membranes and accumulates in the lysosomes of viable cells (Repetto et al. 2008). Toxic substances disrupt this process by impairing cell viability/promoting cell death, which reduces dye uptake. The current NRU assay utilizes a non-epithelial cell line that was not derived from lung tissue and therefore is not associated with respiration. Consequently, the cells lack organ-specific responses relevant to inhalation exposure.

To address this limitation, it is crucial to assess lung–specific cytotoxicity, particularly for airborne chemicals, respiratory drugs, occupational carcinogens, and environmental pollutants. The lung alveoli are primarily lined with two types of epithelial cells: alveolar type I (ATI) and alveolar type II (ATII) cells. ATI cells are responsible for gas exchange, facilitating efficient respiration. In contrast, ATII cells play critical roles in surfactant secretion, immune responses, and maintaining lung architecture and function. Additionally, ATII cells serve as progenitors for ATI cells, contributing to alveolar regeneration and repair following lung injury (Olajuyin et al. 2019; Zacharias et al. 2018), making them essential for toxicological analysis (Cooper et al. 2016; Moreira et al. 2022).

However, conventionally relevant immortalized ATI and ATII cell lines or induced pluripotent stem cell (iPSC)-derived lung cells suitable for standardized lung toxicity assays are currently unavailable. In this context, the human lung adenocarcinoma cell line (A549), which exhibits characteristics of ATII cells, remains the most widely used *in vitro* model for evaluating lung-specific toxicity (Foster et al. 1998; Kramer et al. 2009; Lieber et al. 1976). Given its ATII-like phenotype, the A549 cell line offers greater physiological relevance for evaluating pulmonary toxicity compared to assays utilizing mouse fibroblast 3T3 cells. This enhanced relevance establishes A549 cells as a more effective model for assessing toxicological risks associated with human exposure to airborne substances.

This study aims to develop a novel *in vitro* cytotoxicity test, the A549 cell-based NRU assay (A549-NRU assay), to predict acute inhalation toxicity in rats. We evaluated the cytotoxicity of 49 chemicals, including those associated with occupational and respiratory hazards, and examined the correlation between their physicochemical properties and our *in vitro* LC_50_ values and reported *in vivo* LC_50_ values from 4-hour rat inhalation studies. We also assessed the predictive performance of *in vitro* to *in vivo* extrapolation by evaluating how closely the *in vitro* LC_50_ values aligned with the *in vivo* reference data. Finally, we used our *in vitro* LC_50_ values to estimate LD_50_ values for optimizing starting dose selection for the intra-Tracheal Intra-Pulmonary Spraying (TIPS) method, a simplified alternative to whole-body inhalation studies (Saleh et al. 2025, manuscript under submission).

## Materials and Methods

### 1. Chemicals

The forty-nine chemicals evaluated in this study are listed in Table 1. Neutral red (NR) solution (0.33%) was purchased from Sigma-Aldrich (St. Louis, MO, USA).

**Table 1.**
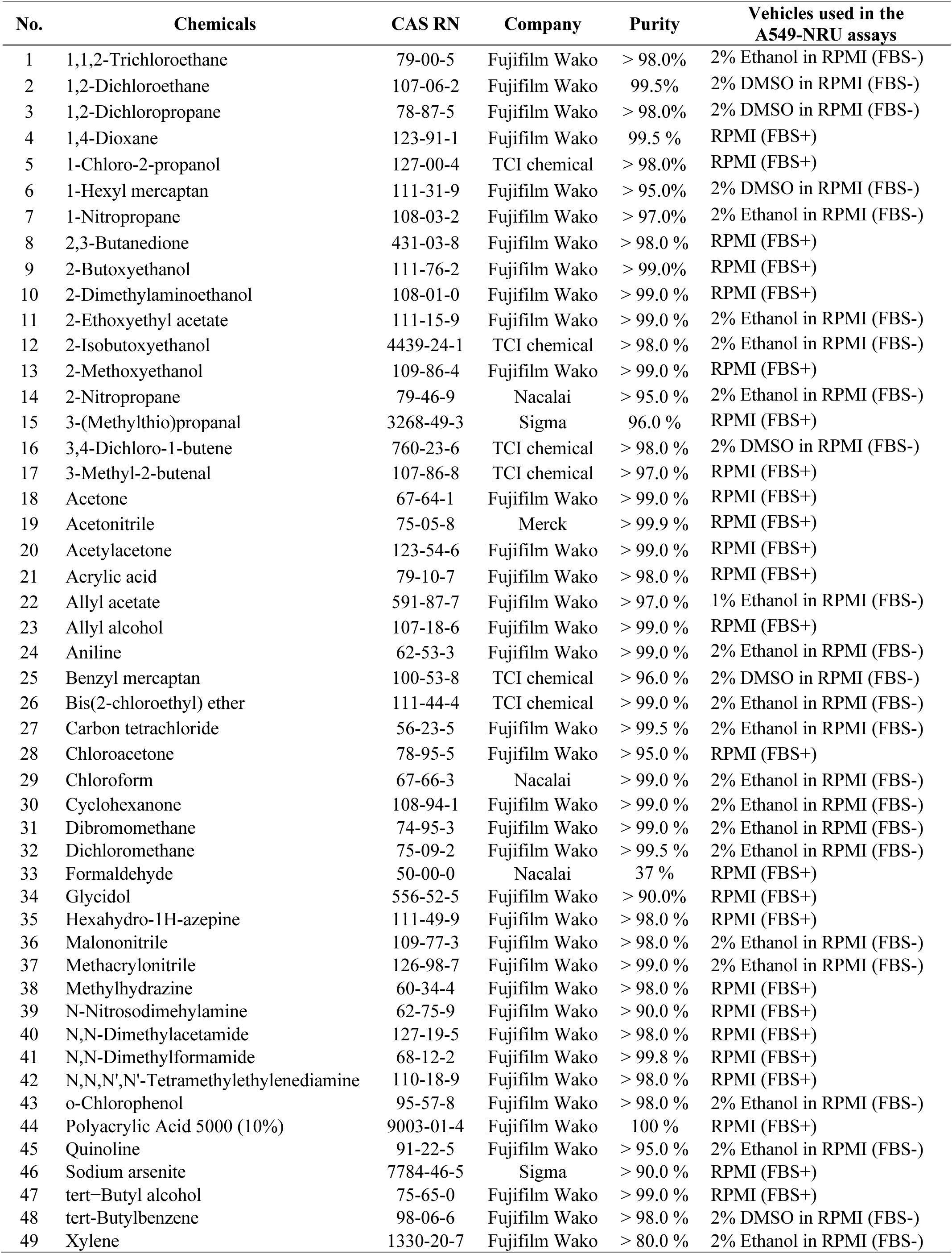
Chemicals evaluated in the A549-NRU assays.

### 2. Cell culture

A549 (RCB3677) cells purchased from RIKEN BioResource Research Center (Ibaraki, Japan) were kindly provided by Dr. Hiroyuki Tsuda (Nagoya City University Graduate School of Medical Sciences). A549 cells were cultured in RPMI-1640 medium with L-glutamine and phenol red (Wako Pure Chemical Industries, Ltd., Osaka, Japan) supplemented with 10% (v/v) fetal bovine serum (FBS) and 1% (v/v) penicillin and streptomycin at 37 °C in an incubator (Sanyo, Japan) with a humidified atmosphere (95% air and 5% CO_2_).

### 3. Neutral red assay

A549 cells were seeded at a density of 2 × 10⁵ cells/well into 6-well polystyrene plates (Corning, New York, USA) or glass plates (Bio Medical Science Inc., Tokyo, Japan), as this density corresponds to the logarithmic growth phase, which is optimal for the present assay. Glass plates were used for 6 chemicals (1,2-dichloropropane, 3,4-dichloro-2-butene, bis(2-chloroethyl) ether, carbon tetrachloride, chloroform, and xylene). These chemicals are reactive with polystyrene, causing it to melt upon contact. The cells were then incubated for 24 h in RPMI-1640 medium with FBS (RPMI(+FBS)). Subsequently, the culture medium was replaced with each test chemical at the designated concentrations, and the cells were incubated at 37 °C for 15 min. Each chemical was evaluated using a wide range of stepwise concentrations in preliminary tests to estimate the approximate LC₅₀ and ensure that it could be determined within a measurable range. LC₅₀ values were calculated using data from at least three concentrations per chemical and were confirmed in at least two independent experiments. Water-soluble chemicals were dissolved in RPMI(+FBS), while water-insoluble chemicals were dissolved in appropriate solvents, up to 2% ethanol (v/v) or 2% DMSO (v/v) in RPMI-1640 medium without FBS (RPMI(-FBS)). Neither 2% ethanol (v/v) nor 2% DMSO (v/v) in RPMI(-FBS) exhibited cytotoxicity in A549 cells following 15-min exposure (data not shown). After incubation, cell viability was examined by the NRU assay following OECD GD 129 and Repetto et al (OECD 2010; Repetto et al. 2008). Briefly, the culture medium containing chemicals was removed and cells were washed twice with PBS. A working NR solution was prepared by diluting a 0.33% (w/v) NR stock solution 1:100 (v/v) in RPMI(+FBS), resulting in a final NR concentration of 0.0033%. The cells were incubated with working NR solution at 37 °C for 3 hours. After incubation, the NR solution was removed, cells were washed twice with PBS, and the intracellular NR dye was extracted using 1ml glacial acetic acid-ethanol (GAE) solution consisting of 1% (v/v) glacial acetic acid, 50% (v/v) ethanol, and 49% (v/v) milli-Q water. Then, 180 µl of the GAE-extract solution was transferred to a 96-well plate and the optical density (OD) was measured at 560 nm (Thermo Fisher Scientific, Massachusetts, USA). All experimental procedures are illustrated in Fig. 1. Cell viability was determined by calculating the ratio of the average OD value of treated wells to the average OD value of control wells. Each condition was tested in two wells per experiment, and the results were confirmed in at least two independent experiments. The mean viability ratios from independent experiments are presented in Table 2. The LC_50_ dose was separately determined for each chemical and calculated by nonlinear regression analysis using GraphPad prism 10 (GraphPad Software, Inc., San Diego, Ca, USA).

**Fig. 1.**
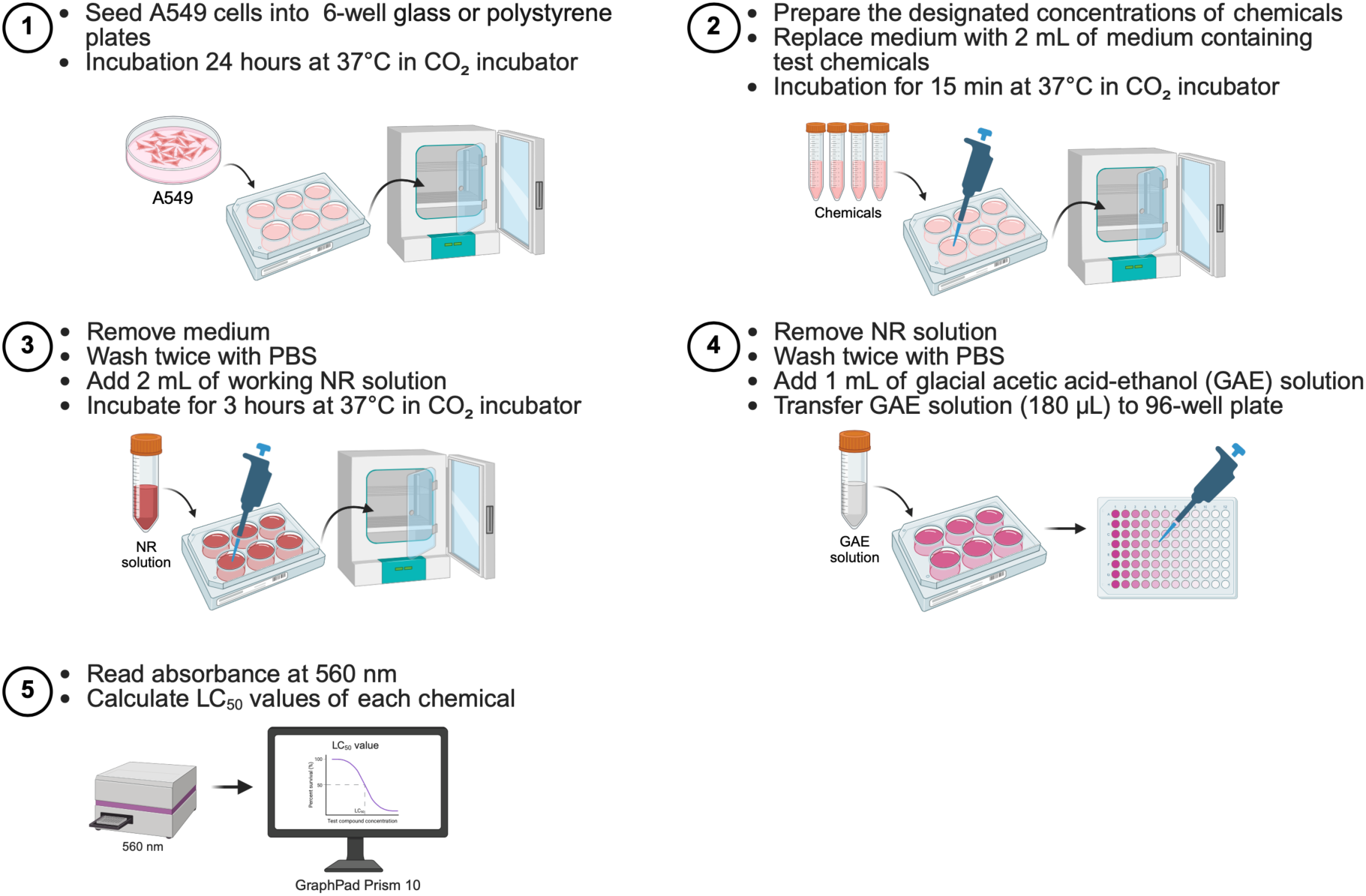
Workflow of the A549-NRU assay. Created in https://BioRender.com.

**Table 2.** Physicochemical properties, LC_50_ values from the A549-NRU assays, and reported LC_50_ values from 4-hour rat inhalation studies for the tested chemicals.

| No. | Chemicals | MW | Density (g/ml) | Log K<sub>ow</sub> | Water soluble | Functional groups | LC<sub>50</sub> from the A549-NRU assays (mg/ml) | LC<sub>50</sub> from 4-hour rat inhalation studies (ppm) | Reference |
| --- | --- | --- | --- | --- | --- | --- | --- | --- | --- |
| 1 | 1,1,2-Trichloroethane | 133.40 | 1.400 | 2.35 | No | Alkyl halide | 2.46 | 2000 | a |
| 2 | 1,2-Dichloroethane | 98.96 | 1.245 | 1.48 | No | Alkyl halide | 7.84 | 1000 | (IARC 1979) |
| 3* | 1,2-Dichloropropane | 112.99 | 1.159 | 1.98 | No | Alkyl halide | 3.33 | 2000 | b |
| 4 | 1,4-Dioxane | 88.11 | 1.034 | -0.42 | Yes | Ether | 117.58 | 9158 | c |
| 5 | 1-Chloro-2-propanol | 94.54 | 1.115 | 0.53 | Yes | Alcohol | 38.97 | 1000 | a |
| 6 | 1-Hexyl mercaptan | 118.24 | 0.840 | 5.35 | No | Thiol | 0.43 | 1080 | c |
| 7 | 1-Nitropropane | 89.09 | 0.996 | 0.87 | No | Nitro | 11.14 | 1512 | b |
| 8 | 2,3-Butanedione | 86.09 | 0.990 | -1.34 | Yes | Ketone | 10.09 | 2250-5200 | b |
| 9 | 2-Butoxyethanol | 118.17 | 0.902 | 0.83 | Yes | Glycol ether | 21.03 | 450 | a |
| 10 | 2-Dimethylaminoethanol | 89.14 | 0.887 | -0.55 | Yes | Amine | 2.59 | 1641 | a |
| 11 | 2-Ethoxyethyl acetate | 132.16 | 0.974 | 0.24 | No | Ester | 39.76 | 2119-3166 | (ECETOC 2005) |
| 12 | 2-Isobutoxyethanol | 118.17 | 0.890 | N/A | No | Glycol Ether | 21.77 | 707-1000 | b |
| 13 | 2-Methoxyethanol | 76.09 | 0.960 | -0.77 | Yes | Glycol ether | 210.53 | 5141 | b |
| 14 | 2-Nitropropane | 89.09 | 0.982 | 0.93 | No | Nitro | 13.02 | 490 | a |
| 15 | 3-(Methylthio)propanal | 104.17 | 1.040 | 0.41 | Yes | Aldehyde | 2.98 | 1036-1105 | a |
| 16* | 3,4-Dichloro-1-butene | 124.99 | 1.153 | 2.37 | No | Alkyl halide | 1.53 | 2100 | c |
| 17 | 3-Methyl-2-butenal | 84.12 | 0.880 | 0.53 | Yes | Aldehyde | 3.90 | 1076 | c |
| 18 | Acetone | 58.08 | 0.785 | -0.24 | Yes | Ketone | 111.87 | 32000 | a |
| 19 | Acetonitrile | 41.05 | 0.787 | -0.54 | Yes | Nitrile | 83.82 | 10679 | c |

**Table 2.** Physicochemical properties, LC<sub>50</sub> values from A549-NRU assays, and reported LC<sub>50</sub> values from 4-hour rat inhalation studies for the tested chemicals (Continued).
| No. | Chemicals | MW | Density (g/ml) | Log K<sub>ow</sub> | Water soluble | Functional groups | LC<sub>50</sub> from the A549-NRU assays (mg/ml) | LC<sub>50</sub> from 4-hour rat inhalation studies (ppm) | Reference |
| --- | --- | --- | --- | --- | --- | --- | --- | --- | --- |
| 20 | Acetylacetone | 100.12 | 0.972 | 0.40 | Yes | Ketone | 15.62 | 1224 | a |
| 21 | Acrylic acid | 72.06 | 1.051 | 0.35 | Yes | Carboxylic acid | 1.88 | 1221 | (IARC 1979) |
| 22 | Allyl acetate | 100.12 | 0.928 | 0.97 | No | Ester | 14.83 | 500 | c |
| 23 | Allyl alcohol | 58.08 | 0.900 | 0.17 | Yes | Alcohol | 39.56 | 165 | a |
| 24 | Aniline | 93.13 | 1.020 | 0.94 | No | Aromatic amine | 6.94 | 250 | (IARC 1982) |
| 25 | Benzyl mercaptan | 124.21 | 1.058 | 2.48 | No | Thiol | 0.55 | 178 | a |
| 26* | Bis(2-chloroethyl) ether | 143.01 | 1.220 | 1.29 | No | Ether | 7.48 | 56 | c |
| 27* | Carbon tetrachloride | 153.82 | 1.594 | 2.64 | No | Alkyl halide | 3.58 | 8000 | a |
| 28 | Chloroacetone | 92.52 | 1.150 | 0.28 | Yes | Ketone | 0.12 | 121 | b |
| 29* | Chloroform | 119.38 | 1.479 | 1.97 | No | Alkyl halide | 5.65 | 2310 | b |
| 30 | Cyclohexanone | 98.14 | 0.942 | 0.81 | No | Ketone | 15.88 | 2450 | b |
| 31 | Dibromomethane | 173.83 | 2.500 | 1.70 | No | Alkyl halide | 9.56 | 3978 | a |
| 32 | Dichloromethane | 84.93 | 1.326 | 1.25 | No | Alkyl halide | 15.90 | 18371 | b |
| 33 | Formaldehyde | 30.03 | 0.815 | 0.35 | Yes | Aldehyde | 3.03 | 480 | b |
| 34 | Glycidol | 74.08 | 1.143 | -0.95 | Yes | Epoxide | 14.02 | 820 | a |
| 35 | Hexahydro-1H-azepine | 99.17 | 0.864 | 1.70 | Yes | Cyclic amine | 1.39 | 604 | a |
| 36 | Malononitrile | 66.06 | 1.190 | -0.60 | No | Nitrile | 30.00 | 52.3-78.6 | a |
| 37 | Methacrylonitrile | 67.09 | 0.800 | 0.68 | No | Nitrile | 17.71 | 328 | a |

**Table 2.** Physicochemical properties, LC<sub>50</sub> values from A549-NRU assays, and reported LC<sub>50</sub> values from 4-hour rat inhalation studies for the tested chemicals (Continued).
| No. | Chemicals | MW | Density (g/ml) | Log K<sub>ow</sub> | Water soluble | Functional groups | LC<sub>50</sub> from the A549-NRU assays (mg/ml) | LC<sub>50</sub> from 4-hour rat inhalation studies (ppm) | Reference |
| --- | --- | --- | --- | --- | --- | --- | --- | --- | --- |
| 38 | Methylhydrazine | 46.07 | 0.870 | -1.05 | Yes | Hydrazine | 4.86 | 78 | a |
| 39 | N-Nitrosodimethylamine | 74.08 | 1.005 | -0.57 | Yes | Nitro | 69.53 | 78 | a |
| 40 | N,N-Dimethylacetamide | 87.12 | 0.937 | -0.77 | Yes | Amide | 141.89 | 1238 | a |
| 41 | N,N-Dimethylformamide | 73.09 | 0.945 | -0.87 | Yes | Amide | 109.18 | 1948 | d |
| 42 | N,N,N',N' Tetramethylethylenediamine | 116.2 | 0.777 | 0.30 | Yes | Amine | 3.81 | 1318 | a |
| 43 | o-Chlorophenol | 128.56 | 1.263 | 2.15 | No | Phenol | 1.24 | 390 | c |
| 44 | Polyacrylic acid 5000 (10%) | 5000 | 0.100 | N/A | Yes | Carboxylic acid | 10.53 | 8 | e |
| 45 | Quinoline | 129.16 | 1.090 | 2.06 | No | Aromatic | 1.53 | N/A | N/A |
| 46 | Sodium arsenite | 129.91 | 1.870 | N/A | Yes | Inorganic | 1.10 | 96 | e |
| 47 | tert-Butyl alcohol | 74.12 | 0.789 | 0.37 | Yes | Alcohol | 45.90 | >10000 | a |
| 48 | tert-Butylbenzene | 134.22 | 0.867 | 4.11 | No | Aromatic | 0.29 | 838 | a |
| 49* | Xylene | 106.17 | 0.870 | 3.20 | No | Aromatic | 0.54 | 5000 | c |

\* Glass plates were used. When the LC<sub>50</sub> value from 4-hour rat inhalation studies is reported as a range, the lower concentration is used for correlation analysis. Information regarding the LD<sub>50</sub> value from 4-hour rat inhalation of chemicals was obtained from the following sources; a: PubChem (https://pubchem.ncbi.nlm.nih.gov), b: National Institute of Technology and Evaluation(NITE, https://www.chem-info.nite.go.jp/chem/english/ghs/ghs_nite_all_fy_e.html), c: Haz-Map (https://haz-map.com/), d: The National Institute for Occupational Safety and Health (NIOSH, https://www.cdc.gov/niosh/rtecs/default.html), and e: Safety Data Sheet (SDS). N/A: Not available.

### 4. Statistical analysis

The Pearson’s correlation coefficient test was performed with GraphPad prism 10, and the significance was determined at *p <* 0.05 vs. the control of each experiment. The criteria for correlation levels are as follows: 0.00 to 0.09 = negligible or very weak correlation; 0.10 to 0.39 = weak correlation; 0.40 to 0.69 = moderate correlation; 0.70 to 0.89 = strong correlation; 0.90 to 1.00 = very strong correlation (Schober et al. 2018).

## Results

### 1. Determination of LC_50_ value and its correlation with the physicochemical properties of tested chemicals in the A549-NRU assays

A total of 49 chemicals were tested, including 24 water-insoluble and 25 water-soluble compounds. The toxicity of all tested chemicals in A549 cells was dose dependent (Supplementary Fig.1). Table 2 summarizes the physicochemical properties, LC_50_ values from the A549-NRU assays, and reported LC_50_ values from 4-hour rat inhalation studies (acute inhalation toxicity test) of all tested chemicals. To analyze the relationship between LC_50_ values from A549-NRU assays and key physicochemical properties, including molecular weight (MW), density, and octanol–water partition coefficient (log K_ow_), Pearson’s correlation analysis was performed, as described below.

#### 1.1 Correlation between LC_50_ values and molecular weight

A correlation analysis was conducted for 48 chemicals, excluding polyacrylic acid (MW = 5000) due to its significantly higher molecular weight compared to the others. The correlation coefficient (Pearson’s r) between LC_50_ values and MW was -0.4206 (*p* = 0.0029), indicating a statistically significant moderate negative correlation (Fig. 2A). This suggests that chemicals with higher MW were associated with lower LC_50_ values, implying greater toxicity.

**Fig. 2.**
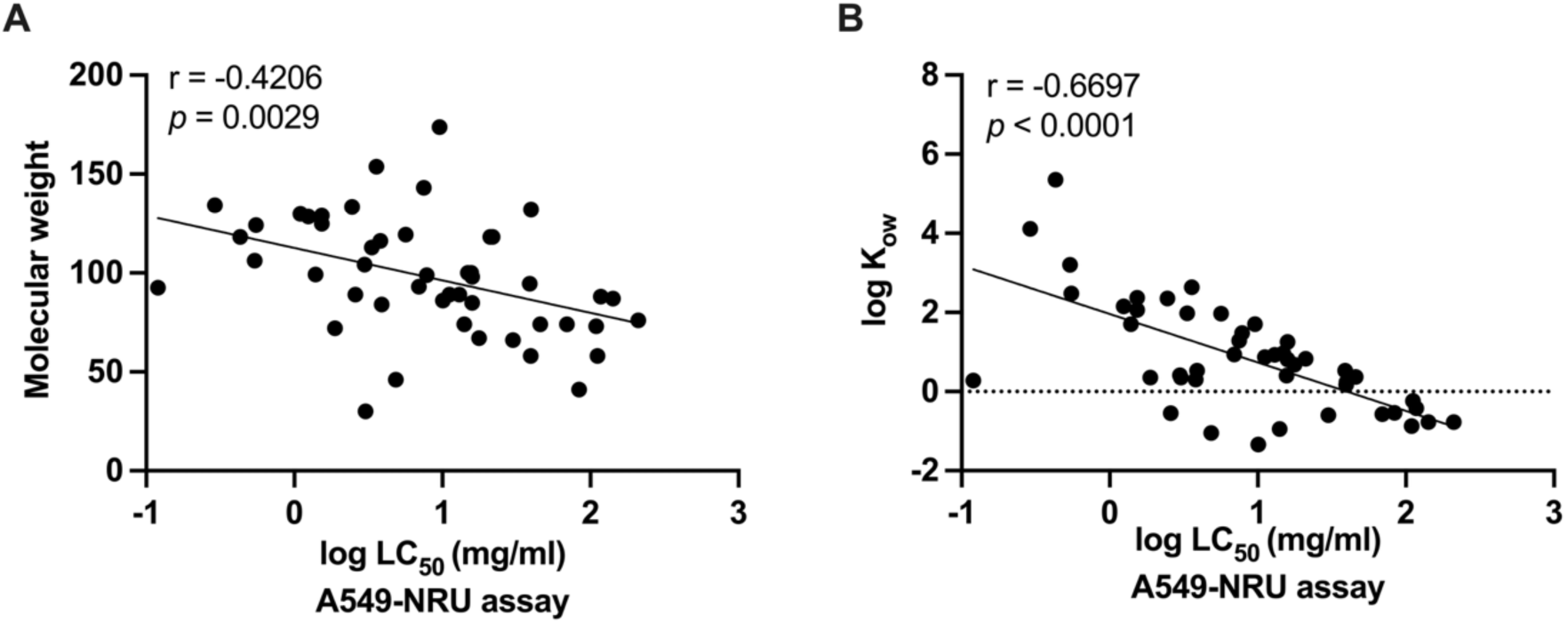
Correlation between LC_50_ values from the A549-NRU assays and molecular weights for 48 tested chemicals (A), correlation with the octanol-water partition coefficient (log K_ow_) for 46 tested chemicals (B).

#### 1.2 Correlation between LC_50_ values and density

The correlation between LC_50_ values and density for all 49 chemicals was weakly negative and not statistically significant (r = -0.1613, *p* = 0.2735), as shown in Supplementary Fig. 2. This suggests that density has no clear association with LC_50_ values, indicating that it is not a suitable parameter for estimating the chemical toxicity in the A549-NRU assay.

#### 1.3 Correlation between LC_50_ values and log K_ow_

Log K_ow_ is a key determinant of a chemical’s distribution between octanol and water, representing its lipophilicity or hydrophilicity. A higher log K_ow_ indicates greater lipophilicity (fat solubility), while a lower log K_ow_ indicates hydrophilicity (water solubility). We assessed the correlation between LC_50_ values and log K_ow_ of 46 chemicals for which the log K_ow_ values are available. The analysis revealed a significantly moderate negative correlation (r = -0.6697, *p* < 0.0001) (Fig. 2B). This indicates that chemicals with higher log K_ow_ (greater lipophilicity) tended to have lower LC_50_ values, suggesting greater toxicity in the A549-NRU assay.

### 2. Correlation analysis of LC_50_ values from the A549-NRU assays and reported LC_50_ values from 4-hour rat inhalation studies based on specific physicochemical properties

To investigate the potential of the A549-NRU assay as a predictive tool for inhalation toxicity, we assessed the correlation between the LC_50_ values obtained from the A549-NRU assays and those reported from 4-hour rat inhalation studies. Among the 49 tested chemicals, quinoline was excluded due to the absence of *in vivo* data, resulting in 48 chemicals included in the analysis (Table 2). Overall, the relationship between LC_50_ values from A549-NRU assays and reported LC_50_ values from 4-hour rat inhalation studies showed a weak positive correlation (r = 0.2504, *p* = 0.0860, Supplementary Fig. 3A). To further elucidate this relationship, 48 chemicals were categorized based on their specific grouping criteria: physicochemical properties including solubility, log K_ow_, and functional groups, which are known to influence chemical behavior and toxicity. This approach aimed to examine whether these properties contribute to variations in the *in vitro-in vivo* correlation.

#### 2.1 Solubility-based analysis

When categorized by solubility, 25 water-soluble chemicals showed a significant moderately positive correlation (r = 0.4632, *p* =0.0197) between the LC_50_ values from the A549-NRU assays and 4-hour rat inhalation studies (Fig. 3A), whereas the 23 water-insoluble chemicals showed a negligible correlation (r = -0.0687, *p* = 0.7553, Supplementary Fig. 3B). These results suggest that LC_50_ values for water-soluble compounds in both assays tended to increase or decrease in parallel, indicating that water solubility may serve as a criterion for predicting LD_50_ values in acute inhalation toxicity tests based on the A549-NRU assay.

**Fig. 3.**
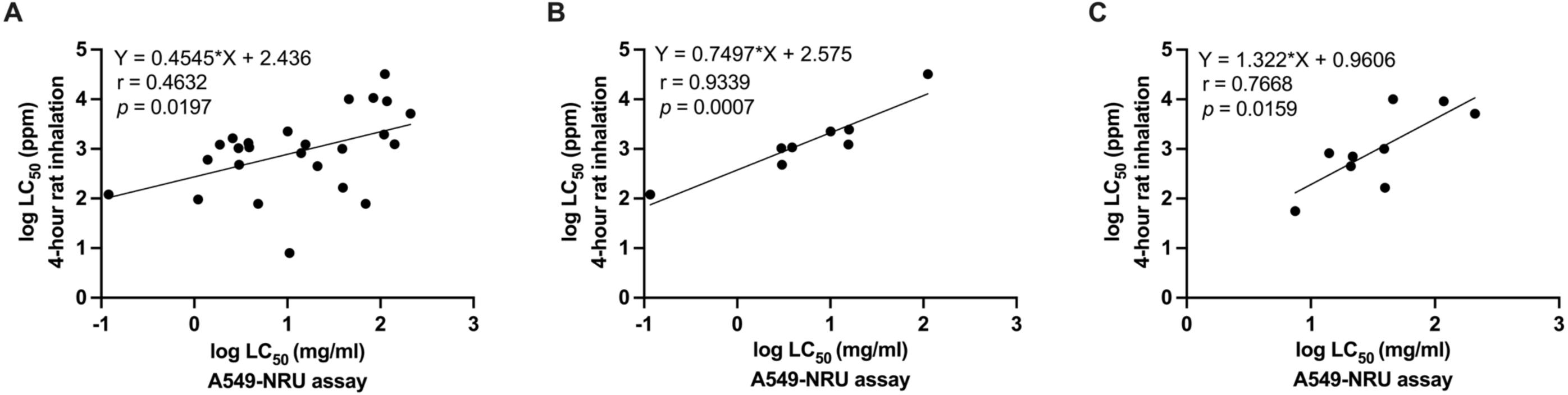
Significant correlations between LC_50_ values from the A549-NRU assays and reported LC_50_ values from 4-hour rat inhalation studies, categorized by water-soluble chemicals (A), chemicals containing aldehyde and ketone groups (B), and chemicals containing alcohol, ether, and epoxide groups (C).

#### 2.2 Log K_ow_-based analysis

45 chemicals were categorized into two groups based on their log K_ow_: negative log K_ow_ (12 chemicals) and positive log K_ow_ (33 chemicals). For the negative log K_ow_ group, a moderate positive correlation (r = 0.4200, *p* = 0.1741) was observed between LC_50_ values from A549-NRU assays and 4-hour rat inhalation studies (Supplementary Fig. 3C). In contrast, the positive log K_ow_ group showed a weak positive correlation (r = 0.1842, *p* = 0.3049, Supplementary Fig. 3D). However, neither correlation was statistically significant, suggesting that log K_ow_ may not be a reliable parameter for estimating starting doses for acute inhalation toxicity with the current set of training chemicals.

#### 2.3 Functional group-based analysis

Based on functional groups, 48 chemicals (all organic compounds) were classified into 10 groups: #1 aldehydes and ketones (8 chemicals), #2 alcohols, ethers, and epoxides (9 chemicals), #3 carboxylic acids and esters (4 chemicals), #4 alkyl halides (8 chemicals), #5 amines and amides (6 chemicals), #6 nitro and nitriles (6 chemicals), #7 aromatics (3 chemicals), #8 thiols (2 chemicals), #9 hydrazine (1 chemical), and #10 phenol (1 chemical). Interestingly, only two groups exhibited statistically significant correlation, group #1, aldehydes and ketones, and group #2, alcohols, ethers, and epoxides. Both groups showed positive correlations between LC_50_ values from the A549-NRU assays and 4-hour rat inhalation studies. Chemicals containing aldehyde and ketone groups exhibited a very strong correlation (r = 0.9339, *p* = 0.0007, Fig. 3B), while those with alcohol, ether, and epoxide groups showed strong correlation (r = 0.7668, *p* = 0.0159, Fig. 3C). These findings indicate that LC_50_ values from A549-NRU assay and 4-hour rat inhalation studies for chemicals containing aldehyde and ketone groups, as well as alcohol, ether, and epoxide groups, tended to align, increasing or decreasing in the same direction. In contrast, the remaining functional groups did not exhibit statistically significant correlations, though some showed negligible, weak, or moderate trends (Supplementary Fig. 4A–4D). Correlation analysis could not be performed for groups #8, #9, or #10 due to the small size in the dataset. In addition, although three chemicals were categorized as aromatics (#7), the analysis could not be conducted because one of them (quinoline) lacked reported LC_50_ data from 4-hour rat inhalation studies.

As described above, among the physicochemical properties, solubility and functional groups emerged as key factors influencing chemical toxicity. Water-soluble chemicals and those containing aldehyde, ketone, alcohol, ether, or epoxide groups may serve as useful criteria for predicting LC_50_ values for inhalation toxicity in rats based on LC_50_ values from A549-NRU assay.

### 3. Predictive regression and concordance analysis between LC_50_ values from the A549-NRU assays and LC_50_ values from 4-hour rat inhalation studies

To evaluate the predictive potential of the A549-NRU assay, group-specific log-linear regression formulas were generated using LC_50_ values from the A549-NRU assays and reported LC_50_ values from 4-hour rat inhalation studies for chemical groups with significant correlations. These formulas were then used to calculate predicted LC_50_ values (ppm) from *in vitro* LC_50_ values (mg/ml), which were subsequently compared with reported *in vivo* LC_50_ values using log_10_ fold change analysis, as summarized in Table 3. The predicted LC_50_ values were considered acceptable if they fell within a ±1 log₁₀ unit (10-fold difference) of the reported values, a commonly applied threshold for extrapolating *in vitro* to *in vivo* doses (Bell et al. 2018; OECD 2010; Spielmann et al. 1999). Using this criterion, 22 of 25 water soluble chemicals, all 8 chemicals containing aldehyde and ketone groups, and all 9 chemicals containing alcohol, ether, and epoxide groups showed predicted LC_50_ values within the acceptable range, supporting the applicability of the regression models. However, as the current dataset contains only the training chemicals used to develop the regression formulas in the predictive model, further validation with a broader set of chemicals is required to confirm their predictive accuracy.

**Table 3.** Comparison of predicted LC_50_ values with reported LC_50_ values from 4-hour rat inhalation toxicity studies for water-soluble chemicals, chemicals containing aldehyde and ketone groups, and chemical containing alcohol, ether, and epoxide groups.

| Chemicals | LC<sub>50</sub> from | 4-hour rat inhalation |  | Log<sub>10</sub> fold | Within ±1<br>log unit<sup>##</sup> |
| --- | --- | --- | --- | --- | --- |
|  | A549-NRU | studies (ppm) |  | change of |  |
|  | assays | Predicted | Reported | predicted LC<sub>50</sub> |  |
|  | (mg/ml) | LC<sub>50</sub><sup>*</sup> | LC<sub>50</sub><sup>**</sup> | to reported LC<sub>50</sub><sup>#</sup> |  |
| Water-soluble chemicals |  |  |  |  |  |
| Formula: Predicted log LC<sub>50</sub> (ppm) = 0.4545 log LC<sub>50</sub> (mg/ml) + 2.436<sup>a</sup> |  |  |  |  |  |
| 1,4-Dioxane | 117.58 | 2382 | 9158 | -0.58 | Yes |
| 1-Chloro-2-propanol | 38.97 | 1442 | 1000 | 0.16 | Yes |
| 2,3-Butanedione | 10.09 | 780 | 2250 | -0.46 | Yes |
| 2-Butoxyethanol | 21.03 | 1089 | 450 | 0.38 | Yes |
| 2-Dimethylaminoethanol | 2.59 | 420 | 1641 | -0.59 | Yes |
| 2-Methoxyethanol | 210.53 | 3104 | 5141 | -0.22 | Yes |
| 3-(Methylthio)propanal | 2.98 | 448 | 1036 | -0.36 | Yes |
| 3-Methyl-2-butenal | 3.90 | 507 | 1076 | -0.33 | Yes |
| Acetone | 111.87 | 2329 | 32000 | -1.14 | No |
| Acetonitrile | 83.82 | 2042 | 10679 | -0.72 | Yes |
| Acetylacetone | 15.62 | 952 | 1224 | -0.11 | Yes |
| Acrylic acid | 1.88 | 363 | 1221 | -0.53 | Yes |
| Allyl alcohol | 39.56 | 1452 | 165 | 0.94 | Yes |
| Chloroacetone | 0.12 | 103 | 121 | -0.07 | Yes |
| Formaldehyde | 3.03 | 452 | 480 | -0.03 | Yes |
| Glycidol | 14.02 | 906 | 820 | 0.04 | Yes |
| Hexahydro-1H-azepine | 1.39 | 317 | 604 | -0.28 | Yes |
| Methylhydrazine | 4.86 | 560 | 78 | 0.86 | Yes |
| N-Nitrosodimethylamine | 69.53 | 1876 | 78 | 1.38 | No |
| N,N-Dimethylacetamide | 141.89 | 2595 | 1238 | 0.32 | Yes |
| N,N-Dimethylformamide | 109.18 | 2303 | 1948 | 0.07 | Yes |
| N,N,N',N'- | 3.81 | 501 | 1318 | -0.42 | Yes |
| Tetramethylethylenediamine |  |  |  |  |  |
| Polyacrylic Acid 5000 | 10.53 | 796 | 8 | 2.00 | No |
| Sodium arsenite | 1.10 | 285 | 96 | 0.47 | Yes |
| t-Butyl alcohol | 45.90 | 1553 | 10000 | -0.81 | Yes |
| Chemicals containing aldehyde and ketone groups |  |  |  |  |  |
| Formula: Predicted log LC<sub>50</sub> (ppm) = 0.7497 log LC<sub>50</sub> (mg/ml) + 2.575<sup>b</sup> |  |  |  |  |  |
| 2,3-Butanedione | 10.09 | 2127 | 2250 | -0.02 | Yes |
| 3-(Methylthio)propanal | 2.98 | 852 | 1036 | -0.09 | Yes |
| 3-Methyl-2-butenal | 3.90 | 1043 | 1076 | -0.01 | Yes |
| Acetone | 111.87 | 12910 | 32000 | -0.39 | Yes |
| Acetylacetone | 15.62 | 2950 | 1224 | 0.38 | Yes |
| Chloroacetone | 0.12 | 75 | 121 | -0.21 | Yes |
| Cyclohexanone | 15.88 | 2987 | 2450 | 0.09 | Yes |
| Formaldehyde | 3.03 | 863 | 480 | 0.25 | Yes |

<sup>\*</sup>Predicted LC<sub>50</sub> (ppm) was calculated using group specific log-linear regression formulas and antilog was applied to obtain LC<sub>50</sub> in ppm. <sup>\*\*</sup>The reported LC<sub>50</sub> values from 4-hour rat inhalation studies (ppm) are from Table 2.
<sup>#</sup>Log<sub>10</sub> fold change = log<sub>10</sub> (predicted LC<sub>50</sub> / reported LC<sub>50</sub>). A value between -1 and +1 indicates that the predicted LC<sub>50</sub> values is within a 10-fold range of the reported value. <sup>##</sup>Indicates whether the log<sub>10</sub> fold change is within ±1 log unit (10-fold range), a commonly accepted threshold for predictive accuracy in toxicity assessment (OECD 2010; Spielmann et al. 1999). <sup>a-c</sup> Regression formulas were derived using Prism 10.

**Table 3.** Comparison of predicted LC<sub>50</sub> values with reported LC<sub>50</sub> values from 4-hour rat inhalation toxicity studies for water-soluble chemicals, chemicals containing aldehyde and ketone groups, and chemical containing alcohol, ether, and epoxide groups (Continued).
| Chemicals | LC<sub>50</sub> from<br>A549-NRU<br>assays<br>(mg/ml) | 4-hour rat inhalation<br>studies (ppm) |  | Log<sub>10</sub> fold<br>change of<br>predicted LC<sub>50</sub><br>to reported LC<sub>50</sub><sup>#</sup> | Within ±1<br>log unit<sup>##</sup> |
| --- | --- | --- | --- | --- | --- |
|  |  | Predicted | Reported |  |  |
|  |  | LC<sub>50</sub><sup>*</sup> | LC<sub>50</sub><sup>**</sup> |  |  |
| **Chemicals containing alcohol, ether and epoxide groups** |  |  |  |  |  |
| Formula: Predicted log LC<sub>50</sub> (ppm) = 1.322 log LC<sub>50</sub> (mg/ml) + 0.9606<sup>c</sup> |  |  |  |  |  |
| 1,4-Dioxane | 117.58 | 4984 | 9158 | -0.26 | Yes |
| 1-Chloro-2-propanol | 38.97 | 1158 | 1000 | 0.06 | Yes |
| 2-Butoxyethanol | 21.03 | 512 | 450 | 0.06 | Yes |
| 2-Isobutoxyethanol | 21.77 | 536 | 707 | -0.12 | Yes |
| 2-Methoxyethanol | 210.53 | 10765 | 5141 | 0.32 | Yes |
| Allyl alcohol | 39.56 | 1181 | 165 | 0.85 | Yes |
| Bis(2-chloroethyl) Ether | 7.48 | 131 | 56 | 0.37 | Yes |
| Glycidol | 14.02 | 300 | 820 | -0.44 | Yes |
| tert-Butyl alcohol | 45.90 | 1437 | 10000 | -0.84 | Yes |

\*Predicted LC<sub>50</sub> (ppm) was calculated using group specific log-linear regression formulas and antilog was applied to obtain LC<sub>50</sub> in ppm. \*\*The reported LC<sub>50</sub> values from 4-hour rat inhalation studies (ppm) are from Table 2.
<sup>#</sup>Log<sub>10</sub> fold change = log<sub>10</sub> (predicted LC<sub>50</sub> / reported LC<sub>50</sub>). A value between -1 and +1 indicates that the predicted LC<sub>50</sub> values is within a 10-fold range of the reported value. <sup>##</sup>Indicates whether the log<sub>10</sub> fold change is within ±1 log unit (10-fold range), a commonly accepted threshold for predictive accuracy in toxicity assessment (OECD 2010; Spielmann et al. 1999). <sup>a-c</sup> Regression formulas were derived using Prism 10.

In addition to the group-specific prediction analysis, overall concordance between LC_50_ values from the A549-NRU assays and those from 4-hour rat inhalation studies (converted from ppm to mg/ml) was examined regardless of physicochemical classifications. This unit conversion enabled direct comparison on the same scale. The log₁₀ fold change between the *in vitro* and converted *in vivo* LC₅₀ values was calculated for each chemical, with concordance defined as falling within a ±1 log₁₀ unit range. As summarized in Table 4, 35 of 48 chemicals met this criterion, further supporting the utility of the A549-NRU assay in initial estimation of acute inhalation toxicity. These findings align with *in vitro* to *in vivo* extrapolation principles, although variability among certain chemicals indicates that additional refinement and validation are required.

**Table 4.** Concordance between LC₅₀ values from A549-NRU assays and converted values from 4-hour rat inhalation studies of all chemicals.

| No. | Chemicals | MW | LC<sub>50</sub> from A549-NRU assays (mg/ml) | Reported LC<sub>50</sub> from 4-hour rat inhalation studies (ppm)* | Estimated LC<sub>50</sub> in lung (mg/ml)** | Log<sub>10</sub> fold change of LC<sub>50</sub> from A549-NRU to reported 4-hour rat inhalation (mg/ml)<sup>#</sup> | Within ±1 log unit<sup>##</sup> |
| --- | --- | --- | --- | --- | --- | --- | --- |
| 1 | 1,1,2-Trichloroethane | 133.40 | 2.46 | 2000 | 41.03 | -1.22 | No |
| 2 | 1,2-Dichloroethane | 98.96 | 7.84 | 1000 | 15.22 | -0.29 | Yes |
| 3 | 1,2-Dichloropropane | 112.99 | 3.33 | 2000 | 34.75 | -1.02 | No |
| 4 | 1,4-Dioxane | 88.11 | 117.58 | 9158 | 124.09 | -0.02 | Yes |
| 5 | 1-Chloro-2-propanol | 94.54 | 38.97 | 1000 | 14.54 | 0.43 | Yes |
| 6 | 1-Hexyl mercaptan | 118.24 | 0.43 | 1080 | 19.64 | -1.66 | No |
| 7 | 1-Nitropropane | 89.09 | 11.14 | 1512 | 20.72 | -0.27 | Yes |
| 8 | 2,3-Butanedione | 86.09 | 10.09 | 2250<sup>a</sup> | 29.79 | -0.47 | Yes |
| 9 | 2-Butoxyethanol | 118.17 | 21.03 | 450 | 8.18 | 0.41 | Yes |
| 10 | 2-Dimethylaminoethanol | 89.14 | 2.59 | 1641 | 22.50 | -0.94 | Yes |
| 11 | 2-Ethoxyethyl acetate | 132.16 | 39.76 | 2119<sup>a</sup> | 43.07 | -0.03 | Yes |
| 12 | 2-Isobutoxyethanol | 118.17 | 21.77 | 707<sup>a</sup> | 12.85 | 0.23 | Yes |
| 13 | 2-Methoxyethanol | 76.09 | 210.53 | 5141 | 60.16 | 0.54 | Yes |
| 14 | 2-Nitropropane | 89.09 | 13.02 | 490 | 6.71 | 0.29 | Yes |
| 15 | 3-(Methylthio)propanal | 104.17 | 2.98 | 1036<sup>a</sup> | 16.60 | -0.75 | Yes |
| 16 | 3,4-Dichloro-1-butene | 124.99 | 1.53 | 2100 | 40.36 | -1.42 | No |
| 17 | 3-Methyl-2-butenal | 84.12 | 3.90 | 1076 | 13.92 | -0.55 | Yes |
| 18 | Acetone | 58.08 | 111.87 | 32000 | 285.82 | -0.41 | Yes |
| 19 | Acetonitrile | 41.05 | 83.82 | 10679 | 67.41 | 0.09 | Yes |
| 20 | Acetylacetone | 100.12 | 15.62 | 1224 | 18.85 | -0.08 | Yes |

**Table 4.** Concordance between LC<sub>50</sub> values from A549-NRU assays and converted values from 4-hour rat inhalation studies of all chemicals (Continued).
| No. | Chemicals | MW | LC<sub>50</sub> from A549-NRU assays (mg/ml) | Reported LC<sub>50</sub> from 4-hour rat inhalation studies (ppm)* | Estimated LC<sub>50</sub> in lung (mg/ml)** | Log<sub>10</sub> fold change of LC<sub>50</sub> from A549-NRU to reported 4-hour rat inhalation (mg/ml)<sup>#</sup> | Within ±1 log unit<sup>##</sup> |
| --- | --- | --- | --- | --- | --- | --- | --- |
| 21 | Acrylic acid | 72.06 | 1.88 | 1221 | 13.53 | -0.86 | Yes |
| 22 | Allyl acetate | 100.12 | 14.83 | 500 | 7.70 | 0.28 | Yes |
| 23 | Allyl alcohol | 58.08 | 39.56 | 165 | 1.47 | 1.43 | No |
| 24 | Aniline | 93.13 | 6.94 | 250 | 3.58 | 0.29 | Yes |
| 25 | Benzyl mercaptan | 124.21 | 0.55 | 178 | 3.40 | -0.79 | Yes |
| 26 | Bis(2-chloroethyl) ether | 143.01 | 7.48 | 56 | 1.23 | 0.78 | Yes |
| 27 | Carbon tetrachloride | 153.82 | 3.58 | 8000 | 189.24 | -1.72 | No |
| 28 | Chloroacetone | 92.52 | 0.12 | 121 | 1.72 | -1.16 | No |
| 29 | Chloroform | 119.38 | 5.65 | 2310 | 42.41 | -0.88 | Yes |
| 30 | Cyclohexanone | 98.14 | 15.88 | 2450 | 36.98 | -0.37 | Yes |
| 31 | Dibromomethane | 173.83 | 9.56 | 3978 | 106.34 | -1.05 | No |
| 32 | Dichloromethane | 84.93 | 15.90 | 18371 | 239.94 | -1.18 | No |
| 33 | Formaldehyde | 30.03 | 3.03 | 480 | 2.22 | 0.14 | Yes |
| 34 | Glycidol | 74.08 | 14.02 | 820 | 9.34 | 0.18 | Yes |
| 35 | Hexahydro-1H-azepine | 99.17 | 1.39 | 604 | 9.21 | -0.82 | Yes |
| 36 | Malononitrile | 66.06 | 30.00 | 52.3<sup>a</sup> | 0.53 | 1.75 | No |
| 37 | Methacrylonitrile | 67.09 | 17.71 | 328 | 3.38 | 0.72 | Yes |
| 38 | Methylhydrazine | 46.07 | 4.86 | 78 | 0.55 | 0.94 | Yes |
| 39 | N-Nitrosodimethylamine | 74.08 | 69.53 | 78 | 0.89 | 1.89 | No |
| 40 | N,N-Dimethylacetamide | 87.12 | 141.89 | 1238 | 16.59 | 0.93 | Yes |

**Table 4.** Concordance between LC<sub>50</sub> values from A549-NRU assays and converted values from 4-hour rat inhalation studies of all chemicals (Continued).
| No. | Chemicals | MW | LC<sub>50</sub> from A549-NRU assays (mg/ml) | Reported LC<sub>50</sub> from 4-hour rat inhalation studies (ppm)* | Estimated LC<sub>50</sub> in lung (mg/ml)** | Log<sub>10</sub> fold change of LC<sub>50</sub> from A549-NRU to reported 4-hour rat inhalation (mg/ml)<sup>#</sup> | Within ±1 log unit<sup>##</sup> |
| --- | --- | --- | --- | --- | --- | --- | --- |
| 41 | N,N-Dimethylformamide | 73.09 | 109.18 | 1948 | 21.90 | 0.70 | Yes |
| 42 | N,N,N',N'-Tetramethylethylenediamine | 116.2 | 3.81 | 1318 | 23.55 | -0.79 | Yes |
| 43 | o-Chlorophenol | 128.56 | 1.24 | 390 | 7.71 | -0.79 | Yes |
| 44 | Polyacrylic Acid 5000 (10%) | 5000 | 10.53 | 8 | 6.15 | 0.23 | Yes |
| 45 | Quinoline | 129.16 | 1.53 | N/A | N/A | - | - |
| 46 | Sodium arsenite | 129.91 | 1.10 | 96 | 1.92 | -0.24 | Yes |
| 47 | tert-Butyl alcohol | 74.12 | 45.90 | 10000 | 113.98 | -0.40 | Yes |
| 48 | tert-Butylbenzene | 134.22 | 0.29 | 838 | 17.30 | -1.78 | No |
| 49 | Xylene | 106.17 | 0.54 | 5000 | 81.64 | -2.18 | No |

\*LC<sub>50</sub> values from 4-hour rat inhalation studies (ppm) are from Table 2. \*\*Estimated LC<sub>50</sub> in the lung (mg/ml) was converted from reported LC<sub>50</sub> (ppm) based on estimated inhaled dose using the formula: $LC_{50} \text{ (mg/ml)} = [(LC_{50} \text{ (ppm)} \times MW \times 0.188 \times 240) / (24.45 \times 1000)] / 12$ , where MW is the molecular weight of each chemical, 0.188 is the estimated respiratory volume (L/min) of a rat with 300 g body weight (Bide et al. 2000), 240 is the exposure duration in minutes, 24.45 is the molar volume of an ideal gas at 25°C and 1 atm, and 12 is the approximated lung vital capacity (ml) of a young rat (300 g) (Hoffman et al. 2014). <sup>#</sup>Log<sub>10</sub> fold change = log<sub>10</sub> (LC<sub>50</sub> from A549-NRU/ LC<sub>50</sub> from 4-hour rat inhalation). A value between -1 and +1 indicates agreement within a 10-fold range. <sup>##</sup>Indicates whether the log<sub>10</sub> fold change is within ±1 log unit (10-fold range), a commonly accepted threshold for predictive accuracy in toxicity assessment (OECD 2010; Spielmann et al. 1999). <sup>a</sup>For LC<sub>50</sub> values reported as a range (Table 2), the lower concentration was used for log<sub>10</sub> fold change analysis. N/A: Not available.

### 4. Comparative analysis of LC_50_ correlations from A549-NRU and 3T3-NRU assays with 4-hour rat inhalation studies

Among the 49 tested chemicals, only 5 chemicals, acetone, acetonitrile, N,N-dimethylformamide, sodium arsenite, and xylene, had reported LC_50_ values from mouse 3T3-NRU assays (Fig. 4A). For correlation analysis, only the water-soluble chemicals were selected due to their significantly moderate positive correlation (Fig. 3A). The correlation between the LC_50_ values from the A549-NRU assays and the reported LC_50_ values from 4-hour rat inhalation studies for these 4 chemicals exhibited a strong positive correlation (r = 0.8879, *p* = 0.1121) (Fig. 4B). This association was notably stronger and more linear compared to the correlation observed between LC_50_ values from the 3T3-NRU assays and reported LC_50_ values from 4-hour rat inhalation studies (r = 0.4524, p = 0.5476) (Fig. 4C).These findings indicate that the A549-NRU assay aligns more closely with rat inhalation toxicity data compared to the 3T3-NRU assay, suggesting that A549 cells are more suitable for predicting inhalation toxicity in rats.

**Fig. 4.**
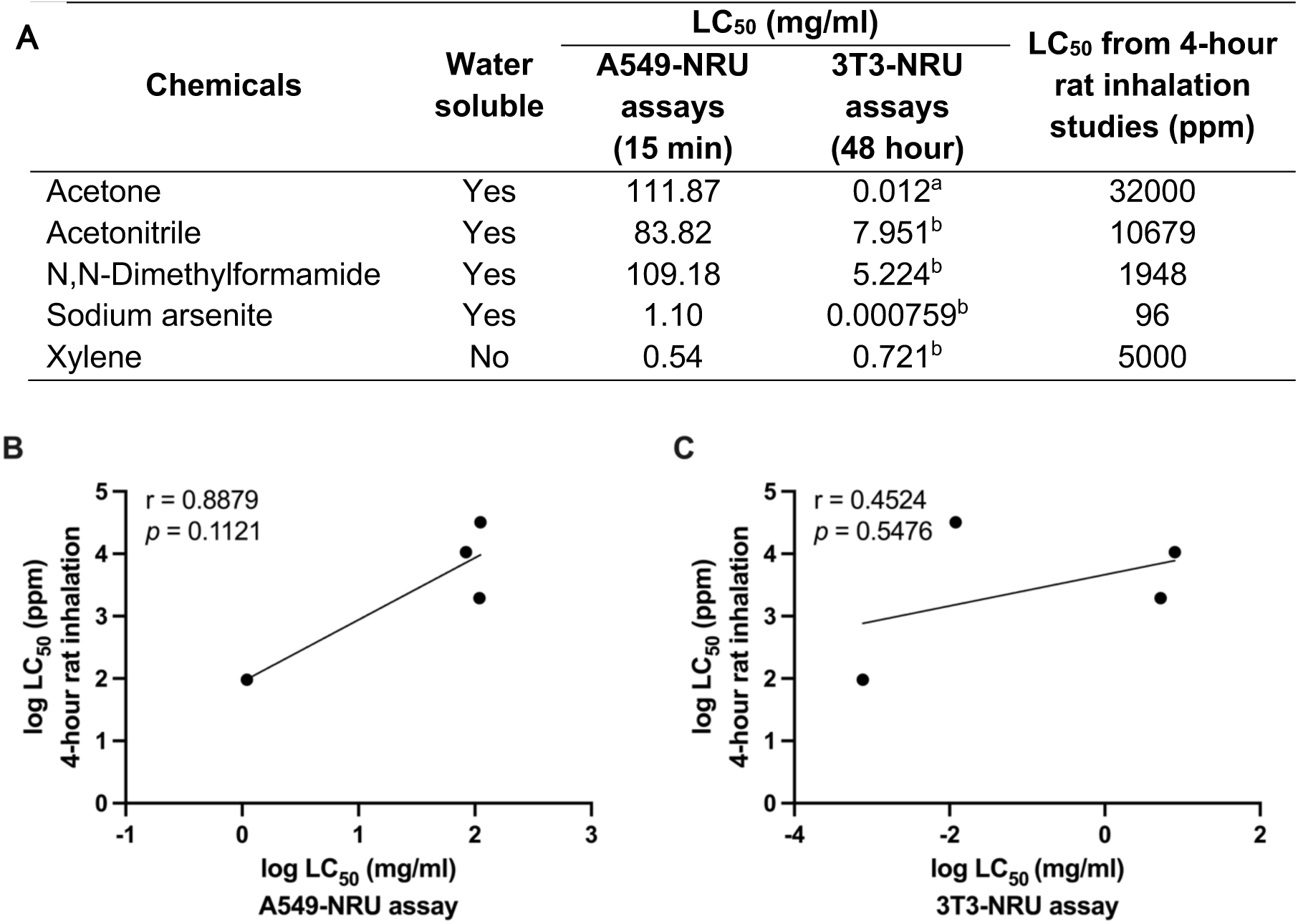
LC_50_ values from A549-NRU and 3T3-NRU assays and reported LC_50_ values from 4-hour rat inhalation studies (A), Correlation between reported LC_50_ values from 4-hour rat inhalation studies and LC_50_ values from A549-NRU assays (B) and LC_50_ values from 3T3-NRU assays (C). Data for the LC_50_ values from 3T3-NRU assays were sourced from ^a^Mannerström et al. (2017), ^b^OECD (2010).

### 5. Estimated LD_50_ values for TIPS based on LC_50_ values from the A549-NRU assay

In addition to using the A549-NRU model to predict 4-hour rat inhalation toxicity, we explored the feasibility of applying this assay to estimate LD_50_ values for the TIPS method through correlation analysis. This approach aims to support the selection of appropriate starting doses and dosing ranges. The estimated LD_50_ values for TIPS, based on the formulation established by Dr. Ogawa and Dr. Tsuda’s groups-collaborative researchers in this study are presented in Table 5. Furthermore, Dr. Ogawa’s group demonstrated that LD_50_ doses calculated from A549-NRU assay LC_50_ values were effective for determining starting doses in TIPS studies (Akane et al. 2025, manuscript under preparation).

**Table 5.**
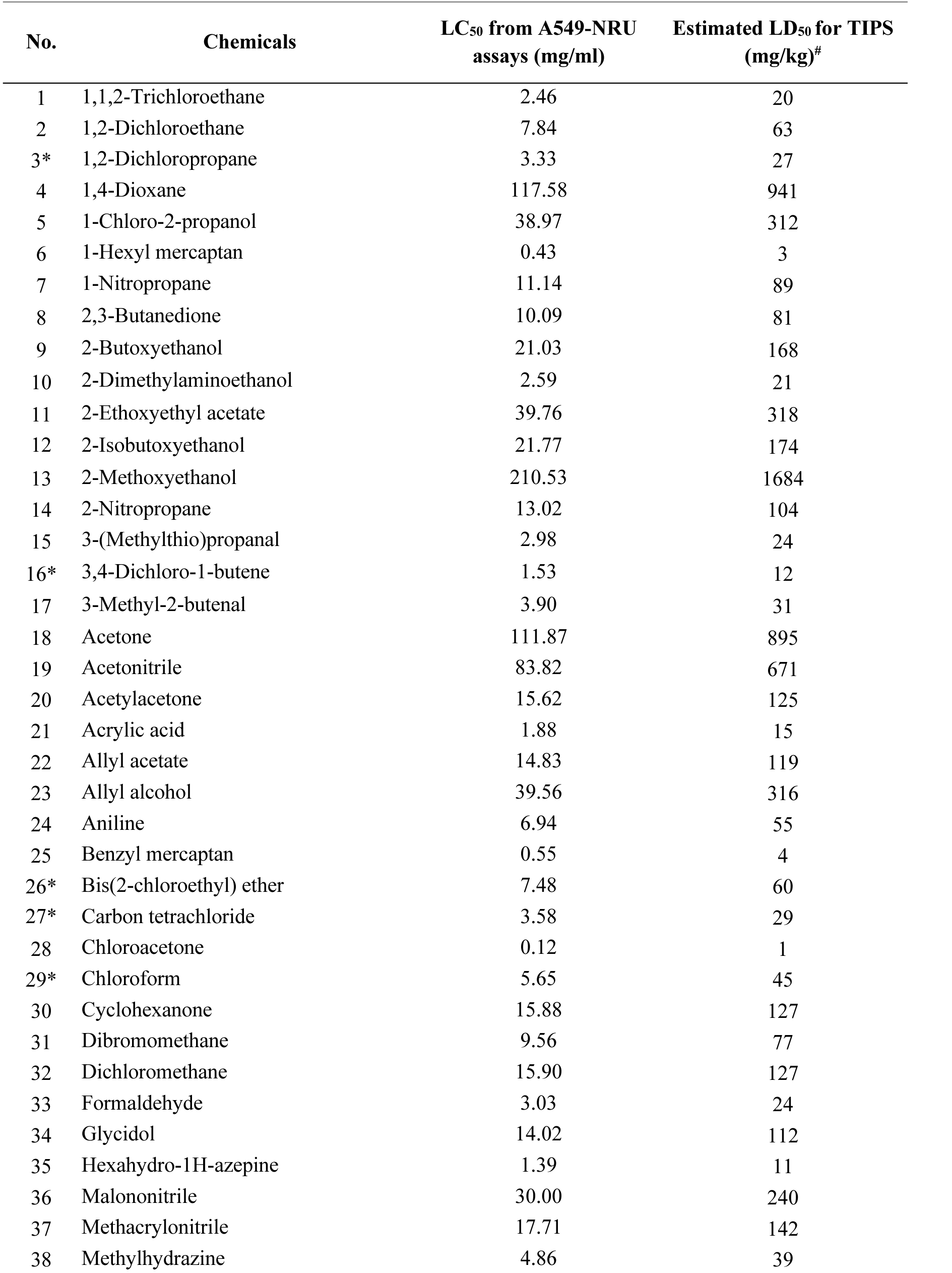
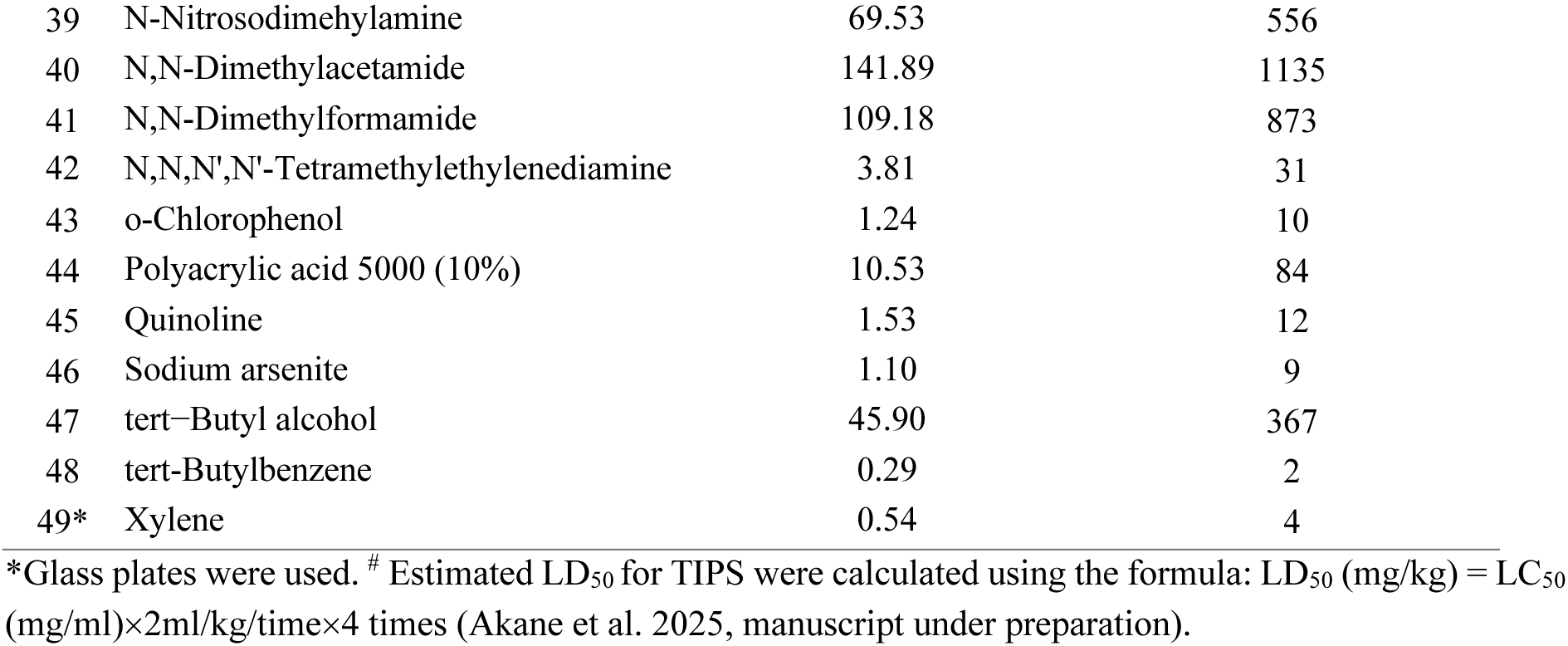
LC_50_ values from the A549-NRU assays and estimated LD_50_ values for TIPS of tested chemicals.

| No. | Chemicals | LC<sub>50</sub> from A549-NRU assays (mg/ml) | Estimated LD<sub>50</sub> for TIPS (mg/kg)<sup>#</sup> |
| --- | --- | --- | --- |
| 1 | 1,1,2-Trichloroethane | 2.46 | 20 |
| 2 | 1,2-Dichloroethane | 7.84 | 63 |
| 3* | 1,2-Dichloropropane | 3.33 | 27 |
| 4 | 1,4-Dioxane | 117.58 | 941 |
| 5 | 1-Chloro-2-propanol | 38.97 | 312 |
| 6 | 1-Hexyl mercaptan | 0.43 | 3 |
| 7 | 1-Nitropropane | 11.14 | 89 |
| 8 | 2,3-Butanedione | 10.09 | 81 |
| 9 | 2-Butoxyethanol | 21.03 | 168 |
| 10 | 2-Dimethylaminoethanol | 2.59 | 21 |
| 11 | 2-Ethoxyethyl acetate | 39.76 | 318 |
| 12 | 2-Isobutoxyethanol | 21.77 | 174 |
| 13 | 2-Methoxyethanol | 210.53 | 1684 |
| 14 | 2-Nitropropane | 13.02 | 104 |
| 15 | 3-(Methylthio)propanal | 2.98 | 24 |
| 16* | 3,4-Dichloro-1-butene | 1.53 | 12 |
| 17 | 3-Methyl-2-butenal | 3.90 | 31 |
| 18 | Acetone | 111.87 | 895 |
| 19 | Acetonitrile | 83.82 | 671 |
| 20 | Acetylacetone | 15.62 | 125 |
| 21 | Acrylic acid | 1.88 | 15 |
| 22 | Allyl acetate | 14.83 | 119 |
| 23 | Allyl alcohol | 39.56 | 316 |
| 24 | Aniline | 6.94 | 55 |
| 25 | Benzyl mercaptan | 0.55 | 4 |
| 26* | Bis(2-chloroethyl) ether | 7.48 | 60 |
| 27* | Carbon tetrachloride | 3.58 | 29 |
| 28 | Chloroacetone | 0.12 | 1 |
| 29* | Chloroform | 5.65 | 45 |
| 30 | Cyclohexanone | 15.88 | 127 |
| 31 | Dibromomethane | 9.56 | 77 |
| 32 | Dichloromethane | 15.90 | 127 |
| 33 | Formaldehyde | 3.03 | 24 |
| 34 | Glycidol | 14.02 | 112 |
| 35 | Hexahydro-1H-azepine | 1.39 | 11 |
| 36 | Malononitrile | 30.00 | 240 |
| 37 | Methacrylonitrile | 17.71 | 142 |
| 38 | Methylhydrazine | 4.86 | 39 |

\*Glass plates were used. <sup>#</sup> Estimated LD<sub>50</sub> for TIPS were calculated using the formula: LD<sub>50</sub> (mg/kg) = LC<sub>50</sub> (mg/ml)×2ml/kg/time×4 times (Akane et al. 2025, manuscript under preparation).

**Table 5.** LC<sub>50</sub> values from A549-NRU assays and estimated LD<sub>50</sub> values for TIPS of tested chemicals (Continued).
| No. | Chemicals | LC<sub>50</sub> from A549-NRU assays (mg/ml) | Estimated LD<sub>50</sub> for TIPS (mg/kg)<sup>#</sup> |
| --- | --- | --- | --- |
| 39 | N-Nitrosodimethylamine | 69.53 | 556 |
| 40 | N,N-Dimethylacetamide | 141.89 | 1135 |
| 41 | N,N-Dimethylformamide | 109.18 | 873 |
| 42 | N,N,N',N'-Tetramethylethylenediamine | 3.81 | 31 |
| 43 | o-Chlorophenol | 1.24 | 10 |
| 44 | Polyacrylic acid 5000 (10%) | 10.53 | 84 |
| 45 | Quinoline | 1.53 | 12 |
| 46 | Sodium arsenite | 1.10 | 9 |
| 47 | tert-Butyl alcohol | 45.90 | 367 |
| 48 | tert-Butylbenzene | 0.29 | 2 |
| 49* | Xylene | 0.54 | 4 |

\*Glass plates were used. <sup>#</sup> Estimated LD<sub>50</sub> for TIPS were calculated using the formula: LD<sub>50</sub> (mg/kg) = LC<sub>50</sub> (mg/ml)×2ml/kg/time×4 times (Akane et al. 2025, manuscript under preparation).

## Discussion

Inhalation toxicities of chemicals are, in principle, classified based on LC_50_ values obtained from acute inhalation toxicity studies that typically require a large number of animals. However, there is increasing demand for alternative evaluation methods that utilize *in vitro* assays to assess toxicity. To address this need, we partially modified the 3T3-NRU cytotoxicity assay described in OECD Guidance Document 129 (OECD 2010). We replaced the 3T3 fibroblast cell line with the human lung adenocarcinoma cell line A549, which better reflects lung-specific functions and offers a more appropriate model for assessing inhalation hazards. Using the optimized A549-NRU assay, we calculated LC_50_ values for 49 substances, demonstrating its potential utility for evaluating *in vivo* inhalation toxicity.

Our A549-NRU assay incorporates key modifications. First, reducing the incubation time to 15 minutes enabled rapid and efficient screening of multiple chemicals, thereby enhancing throughput. Additionally, to address the issue of chemical-induced degradation of polystyrene plates, we employed 6-well glass plates instead of polystyrene plates when testing chemicals that reacted with polystyrene. This approach not only prevented plate deformation but also maintained consistent results, making glass plates a practical alternative for handling a wide range of reactive test chemicals. To our knowledge, this is the first study to use glass plates in this context, and their performance was comparable to that of polystyrene plates.

Physicochemical properties are crucial in determining the cellular toxicity of substances by influencing their interactions with various cellular components (Brinkmann et al. 2014; Kramer et al. 2009). Among the parameters examined, MW and log K_ow_ exhibited significant negative correlations with LC_50_ values in the A549-NRU assay. These inverse relationships imply that chemicals with lower MW and log K_ow_ tend to be less toxic. Compounds with lower MW and smaller sizes can penetrate cellular membranes more efficiently through passive diffusion, reaching intracellular targets more rapidly and potentially causing toxic effects. However, these compounds are also typically excreted from cells more rapidly and efficiently than larger molecules, thereby limiting their bioaccumulation and reducing their toxicity potential (Yang and Hinner 2015). Consistent with this, our study demonstrates that chemicals with low MW exhibited higher LC₅₀ values, indicating lower toxicity, which may be attributed to more rapid elimination from cells.

Chemicals with higher log K_ow_ (greater lipophilicity) typically exhibit lower LC_50_ values, indicating increased toxicity. Log K_ow_ is a widely used parameter in toxicology and pharmacology as a model for chemical partitioning between blood and lipid membranes and serves as a key predictor of membrane permeability (Lee and Lin 2014). The increased toxicity arises from their reduced water solubility, enhanced membrane permeability, and greater intracellular bioaccumulation within lipid-rich cellular environments. These properties enable chemicals to efficiently penetrate cellular membranes, disrupt biochemical processes and induce cytotoxic effects at relatively lower concentrations (Chmiel et al. 2019; Yukawa and Naven 2020). Although some chemicals with low log K_ow_ also exhibited high toxicity, most demonstrated negative correlations between LC_50_ and log K_ow_. The significant negative correlations of MW and log K_ow_ with LC_50_ values in the A549-NRU assay suggest their relevance as predictors of chemical toxicity.

To further assess the relevance of our *in vitro* findings, we grouped the tested chemicals based on the presence of functional groups. Structural complexity or the presence of toxicophores frequently involve functional groups that enhance reactivity with cellular macromolecules, which can further enhance toxicity through enzyme inhibition, oxidative stress, and genotoxicity (Singh et al. 2016; Webel et al. 2020). Among the different groups, only two groups, one including aldehyde and ketone groups and another including alcohol, ether, and epoxide groups, exhibited strong *in vitro*–*in vivo* toxicity correlations. These structures are known to influence chemical reactivity and interactions with cellular macromolecules, which may explain their consistent toxicity profiles across both *in vitro* and *in vivo* systems. For instance, the carbonyl groups (C=O) in aldehydes and ketones are electrophilic and readily form covalent bonds with nucleophilic sites on proteins and DNA, potentially disrupting essential cellular functions (LoPachin and Gavin 2014; Schultz and Yarbrough 2004). Similarly, alcohols, ethers, and epoxides can participate in hydrogen bonding with some amino acid side chains, resulting in disruption of protein structure and function (Alabugin et al. 2021; Oroguchi and Nakasako 2017). It is possible that these functional groups interact with pulmonary cells in a similar manner both *in vitro* and *in vivo*, thereby contributing to their predictive reliability.

Water-soluble chemicals also showed positive correlation between *in vitro* and *in vivo* LC_50_ values. This suggests that these chemicals are soluble to a similar extend in the fluid lining the lung epithelium and the media used for the A549-NRU assay.

On the other hand, water-insoluble compounds and chemicals categorized by groupings of functional groups 3-6 exhibited statistically non-significant correlations between A549-NRU assay LC₅₀ values and 4-hour rat inhalation studies. In addition, while there was a significant negative correlation of log K_ow_ with A549-NRU LC_50_ values, chemicals with low log K_ow_ values and negative log K_ow_ values did not correlate well with A549-NRU assay LC₅₀ values, see Figure 2B. Moreover, the correlation of LC_50_ values from A549-NRU assays and 4-hour rat inhalation studies was not significant for chemicals with negative log K_ow_ values (Supplementary Fig. 3C). Notably however, the correlation of LC_50_ values from A549-NRU assays and 4-hour rat inhalation studies was also not significant for chemicals with positive log K_ow_ values (Supplementary Fig. 3D). These inconsistencies may be attributed to physiological differences between *in vitro* and *in vivo* systems, especially in metabolism, bioavailability, and accumulation. Although log K_ow_ clearly influences chemical toxicity in *in vitro* systems, it may not serve as an effective predictor of *in vivo* acute inhalation toxicity. Overall, these findings highlight the complexity of extrapolating *in vitro* results to *in vivo* outcomes and the importance of considering a variety of parameters to improve predictive accuracy. This variability indicates that further studies are necessary to characterize these effects.

To examine the predictive potential of the A549-NRU assay in estimating *in vivo* inhalation toxicity, we applied a commonly accepted threshold for IVIVE. *In vitro* LC_50_ values from the A549-NRU assay were considered concordant with *in vivo* LC_50_ values if they were within ±1 log₁₀ unit (within a 10-fold difference range), which is an acceptable criteria used in IVIVE studies (Bell et al. 2018; Krewski et al. 2020; OECD 2010; Spielmann et al. 1999). In this context, the application of IVIVE provides a feasible strategy for estimating *in vivo* toxicity values, thereby supporting dose selection and reducing reliance on animal testing. We performed two types of concordance evaluations: one using regression-based prediction of *in vivo* LC_50_ values derived from A549-NRU data, and the other by directly comparing *in vitro* and *in vivo* LC_50_ values after unit conversion. Both approaches yielded comparable results, with a substantial proportion of chemicals falling within a 10-fold difference range. These findings support the predictive potential of the A549-NRU assay for estimating acute inhalation toxicity. However, validation using a broader set of chemicals is required to further refine and improve the accuracy of the predictive model.

In addition to its role in predicting acute inhalation toxicity in rats, the A549-NRU assay also showed potential for estimating LD_50_ values for chemicals administered by the TIPS method, an alternative model for determining respiratory toxicity (Saleh et al. manuscript submitted). Our collaborative research groups (Ogawa K) have already applied LC_50_ values from the A549-NRU assay to calculate LD_50_ doses for TIPS administration (Akane et al. 2025, manuscript under preparation). These values served as a useful reference to estimate starting doses, and a manuscript is currently under preparation.

Mouse 3T3 fibroblast cells are commonly used in the NRU assay and recommended by European Centre for the Validation of Alternative Methods (ECVAM) for regulatory use under certain conditions (OECD 2010). However, the comparison of LC_50_ values from the A549-NRU and 3T3-NRU assays with LC_50_ values from 4-hour rat inhalation reveals a difference in predictive capacity between these models. Our findings demonstrate that LC_50_ values from A549 cells exhibit a much stronger correlation with LC_50_ values from 4-hour rat inhalation exposure than those from mouse 3T3 cells. The greater correlation observed with A549 cells, which originate from human alveolar epithelial type II cells, may be attributed to their tissue-specific physiological relevance to the respiratory system (Foster et al. 1998; Lieber et al. 1976). Furthermore, A549 cells express several cytochrome P450 enzymes, which are important for the metabolic activation and inactivation of many hazardous substances, as these chemicals require enzymatic biotransformation to reactive forms (toxic forms) (Hukkanen et al. 2000). In contrast, 3T3 cells, derived from murine fibroblasts (OECD 2019), are less representative of lung tissue, with differences in cellular uptake, metabolic capacity, and tissue-specific responses possibly explaining their weaker correlation with *in vivo* LC_50_ results. Taken together, these factors suggest that the A549-NRU assay provides a more reliable estimate of acute inhalation toxicity compared to mouse 3T3 fibroblasts, thereby improving the predictability of animal toxicity in the NRU assay.

In conclusion, these findings highlight the importance of considering the physicochemical properties of chemicals when assessing their toxicity. The A549-NRU assay demonstrates high potential for predicting acute inhalation toxicity in the rat and estimating the starting dose for the TIPS method, particularly for water-soluble chemicals and those containing aldehyde ketone, alcohol, ether, and epoxide groups. Overall, A549 cells appear to be a more appropriate choice than 3T3 cells for estimating inhalation toxicity in rats, providing valuable insights for inhalation toxicology. In the future, it will be necessary to further expand the versatility and improve the predictive accuracy of the A549-NRU assay through validation with a broader range of substances and the accumulation of additional data.

## Acknowledgements

This work was supported by Health and Labour Sciences Research Grants from the Ministry of Health, Labour and Welfare of Japan (22KD1003).

## Data Availability Statement

Data will be made available on request.

