## Supplementary Fig.1-4 for "Developing a Novel *In Vitro* Toxicity Assay for Predicting Inhalation Toxicity in Rats"

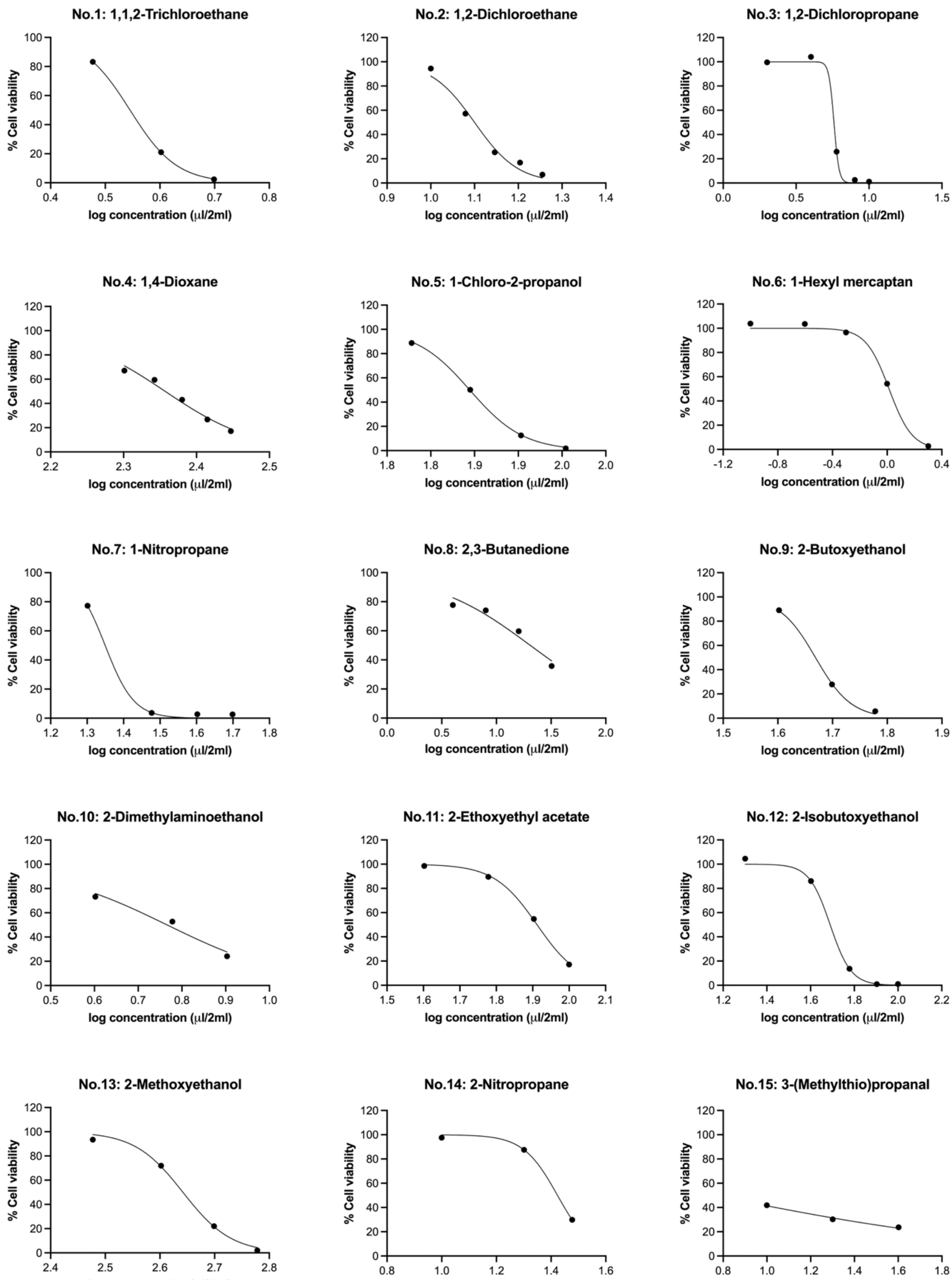


**Supplementary Fig. 1.** Dose-response curves of all tested chemicals in the A549-NRU assays.

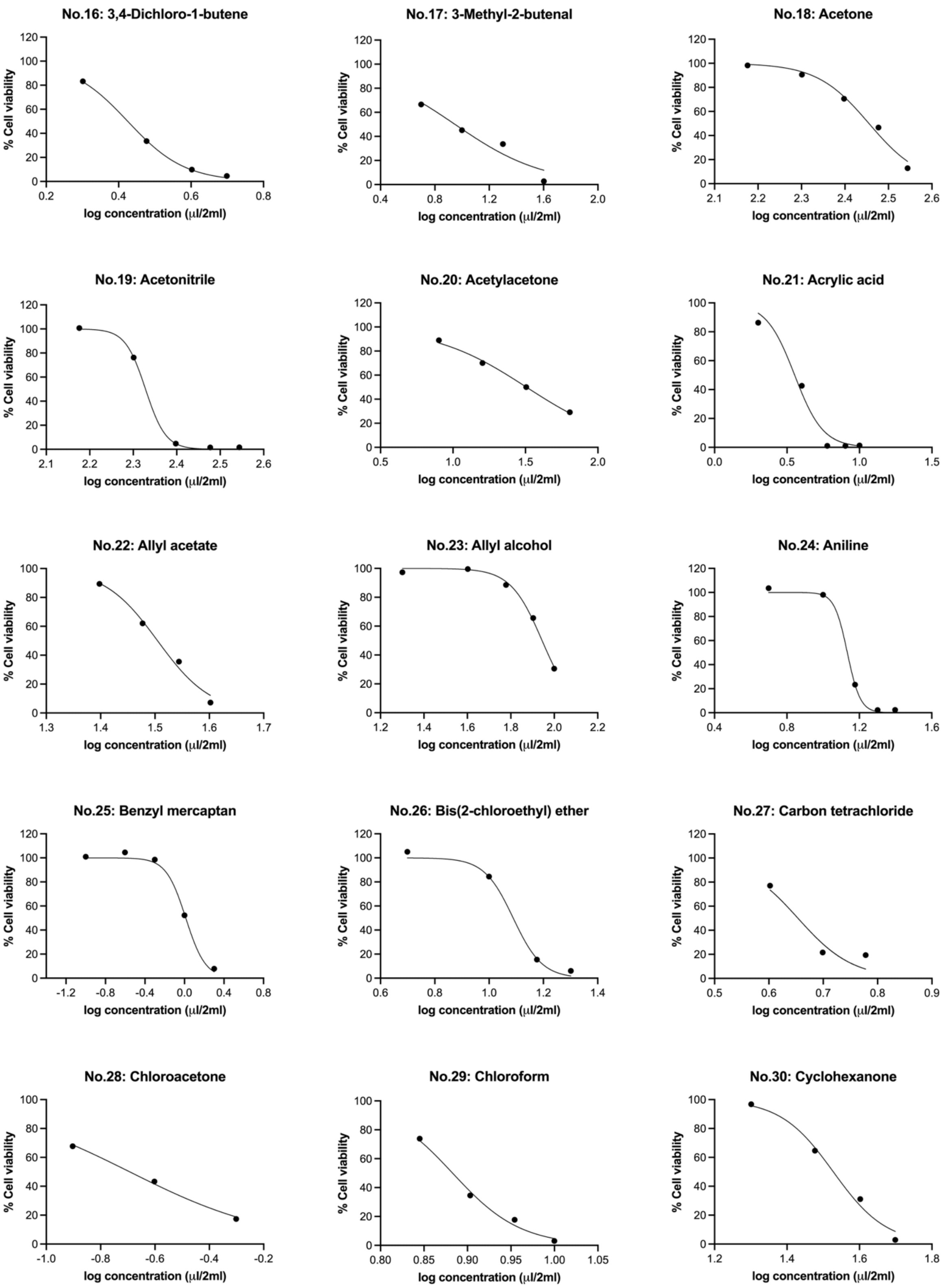


**Supplementary Fig. 1.** Dose-response curves of all tested chemicals in the A549-NRU assays (Continued).

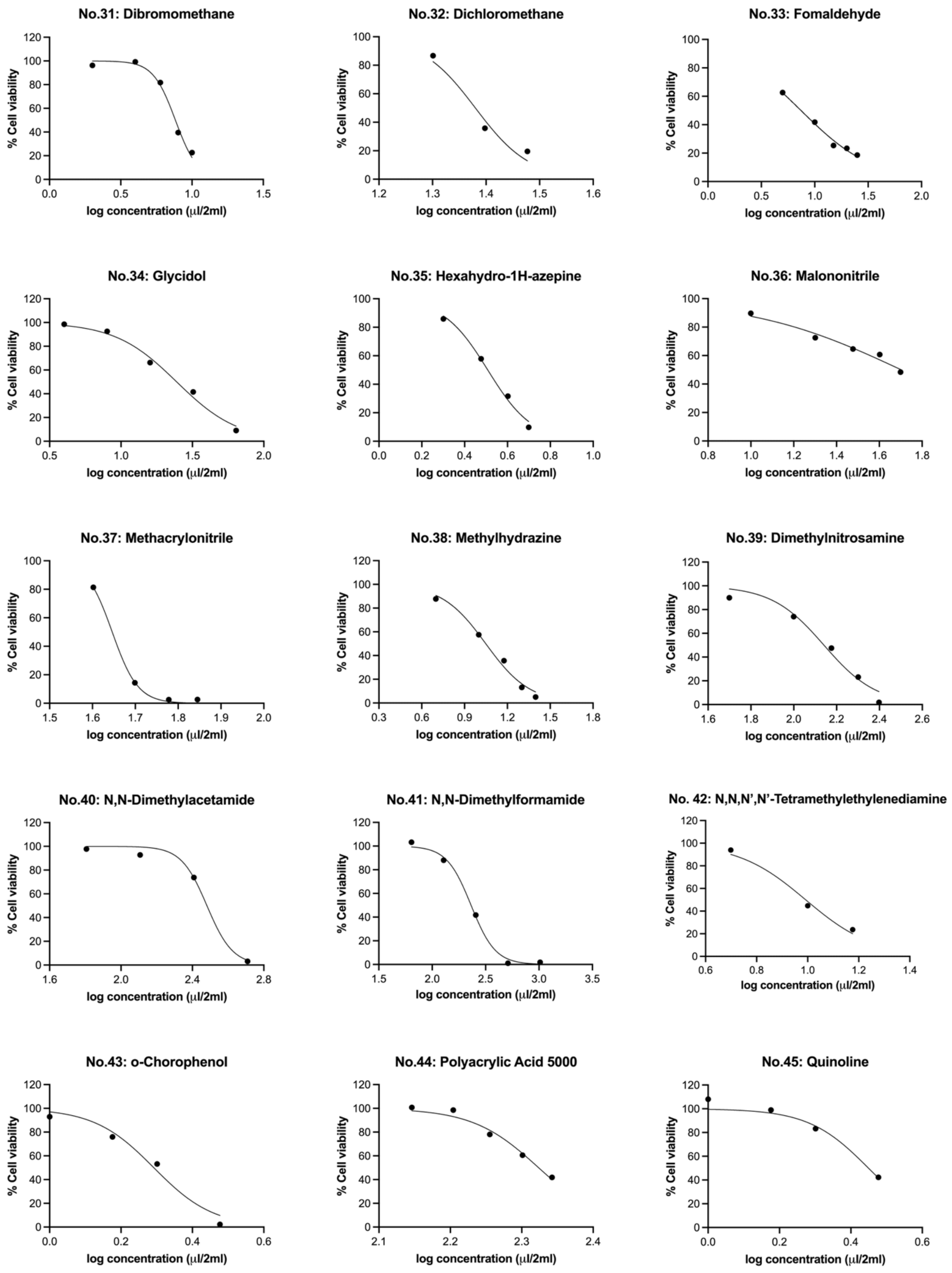


**Supplementary Fig. 1.** Dose-response curves of all tested chemicals in the A549-NRU assays (Continued).

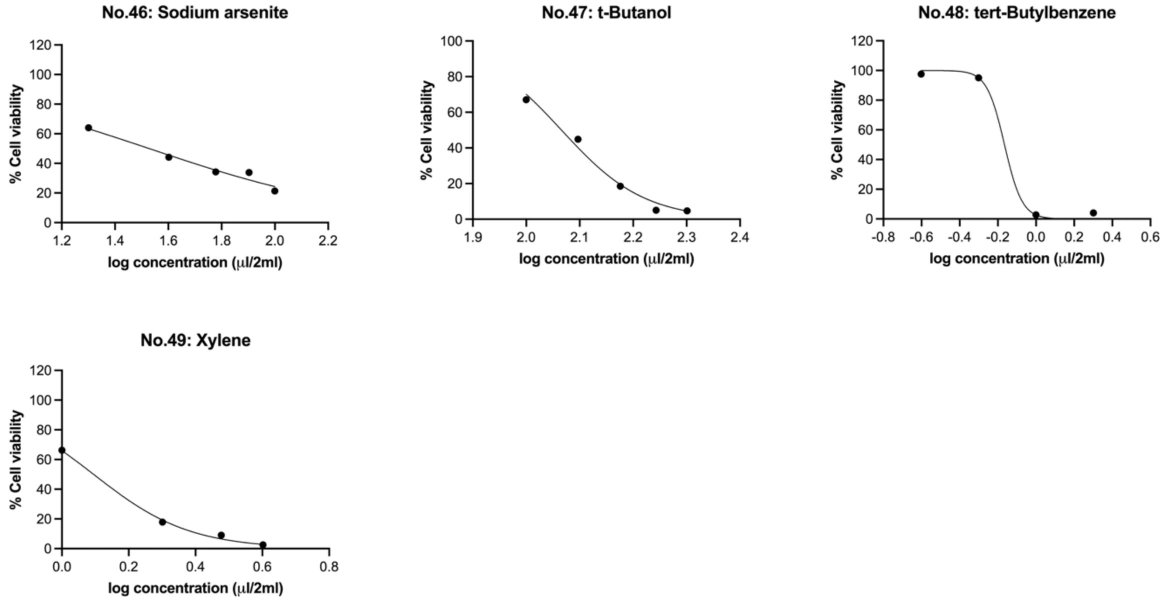


**Supplementary Fig. 1.** Dose-response curves of all tested chemicals in the A549-NRU assays (Continued).

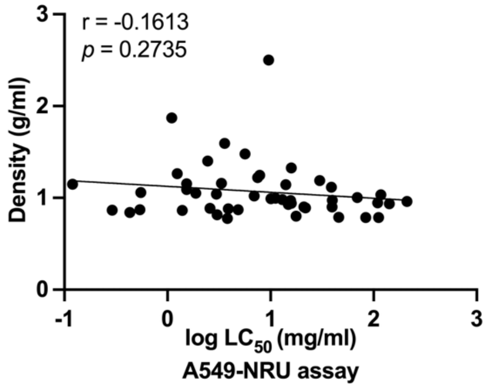


**Supplementary Fig. 2.** The correlation between log $LC_{50}$ values form A549-NRU assays and density of all chemicals.

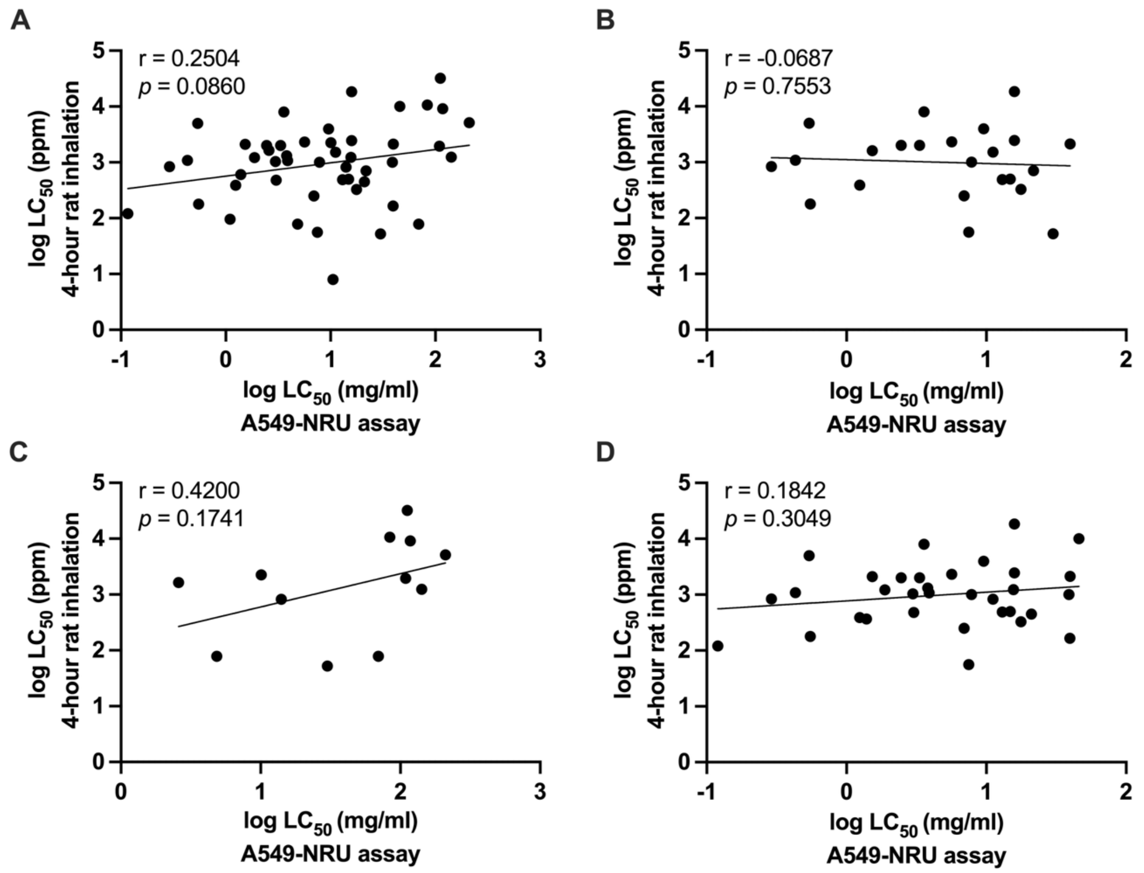


**Supplementary Fig. 3.** The correlation between $LC_{50}$ values from A549-NRU assays and reported $LC_{50}$ values from 4-hour rat inhalation studies of all chemicals (A), chemicals with water-insoluble (B), negative log $K_{ow}$ (C), and positive log $K_{ow}$ (D).

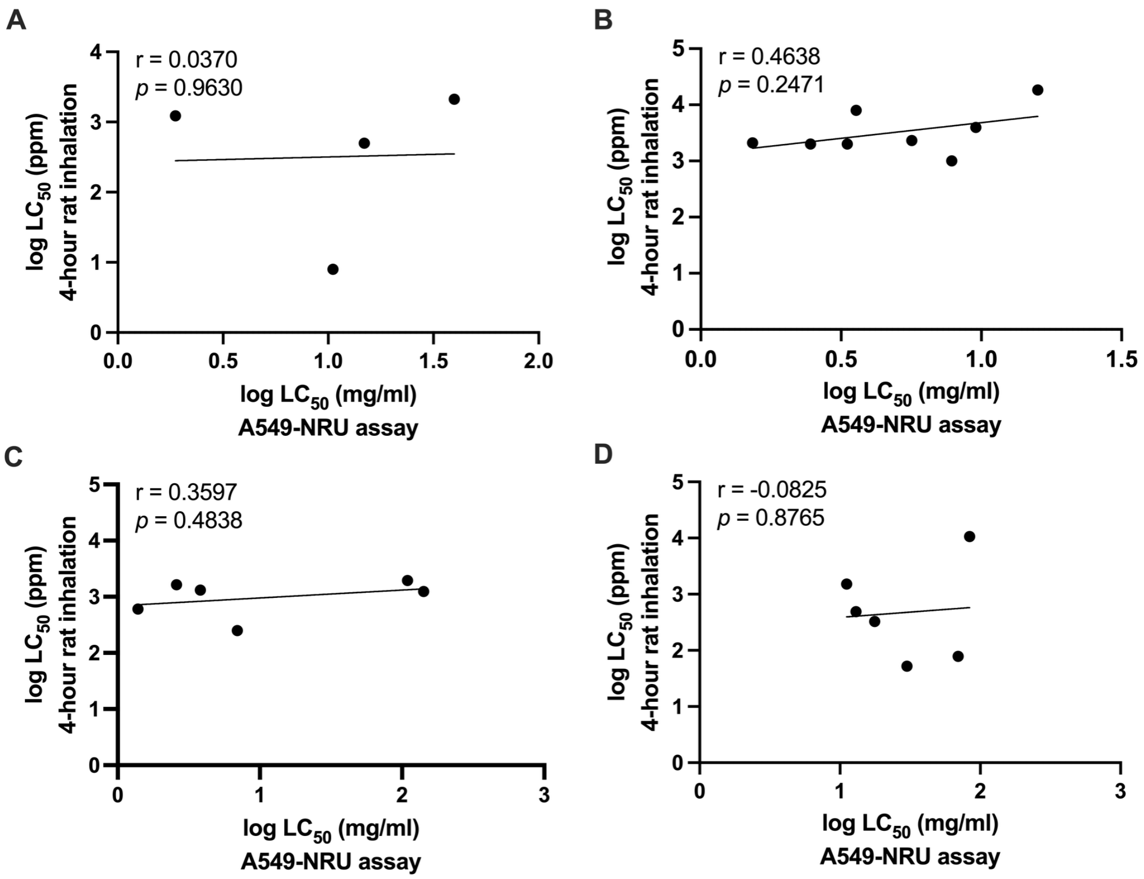


**Supplementary Fig. 4.** The correlation between $LC_{50}$ values from A549-NRU assays and reported $LC_{50}$ values from 4-hour rat inhalation studies categorized by their functional groups: carboxylic acid and ester (A), alkyl halide (B), amine and amide (C), and nitro and nitrile (D).
